# Identification and biosynthesis of xildivaline, a novel and widespread peptide deformylase inhibitor from Gammaproteobacteria

**DOI:** 10.1101/2025.06.07.658423

**Authors:** Alexander Rill, Margaretha Westphalen, Morgan Lamberioux, Janine Chekaiban, Chloé Janin, Didier Mazel, Michael Groll, Eva M. Huber, Helge B. Bode

**Affiliations:** Department of Natural Products in Organismic Interactions, Max Planck Institute for Terrestrial Microbiology, 35043 Marburg, Germany; Molecular Biotechnology, Department of Biosciences, Goethe University Frankfurt, 60438 Frankfurt am Main, Germany; Institut Pasteur, Université Paris Cité, Unité Plasticité du Génome Bactérien, CNRS UMR3525, Paris, France; Collège Doctoral, Sorbonne Université, Paris, France; Technical University of Munich, TUM School of Natural Sciences, Department of Bioscience, Center for Protein Assemblies, 85748 Garching, Germany; Chemical Biology, Department of Chemistry, Philipps University of Marburg, 35043 Marburg, Germany; Center for Synthetic Microbiology (SYNMIKRO), Philipps University of Marburg, 35043 Marburg, Germany

## Abstract

*Xenorhabdus* strains, Gram-negative bacteria pathogenic to insects and symbionts to nematodes of the genus *Steinernema* are prolific producers of various natural products. Here we describe the *xisABCDE* biosynthesis gene cluster from *Xenorhabdus hominickii* responsible for the production of xildivalines. These non-ribosomal peptide and polyketide hybrids act as peptide deformylase inhibitor (PDI) and occur also in other Gammaproteobacteria, especially *Vibrio*. Their structure and biosynthesis were fully elucidated despite their instability, highlighting a rare *trans*-methylation of their N-terminus. Subsequently, the structure of the responsible methyltransferase XisE and the peptide deformylase XisD, serving as resistance mechanism, were elucidated by X-ray crystallography, allowing insights into the function and the mode of action of this novel class of PDIs.

## Introduction

While microbial natural products are important drugs applied especially as antibiotics, the increase in antimicrobial resistance (AMR) of various human pathogens renders current antibiotics useless. Multiple efforts from various disciplines and stakeholders are needed to fuel the ever-increasing demand to treat such otherwise life-threatening infections.^[1]^ One such approach could be the identification of novel microbial natural products (NPs), which is facilitated by excellent bioinformatic tools,^[2]^ which even allow to prioritize biosynthetic gene clusters (BGCs) based on encoding resistance genes pointing towards a desired bioactivity.^[3,4]^ Such approaches might speed up drug discovery even more in the future when artificial intelligence can be applied to identify novel natural product structures from complex mass spectrometry data, the BGCs producing them and ideally also predict their biological targets.^[5]^ A big-data independent and probably more classical approach is to search natural products at places where they are needed, thus applying microbial ecology knowledge. This has been tremendously successful even when we often do not fully understand the ecology behind the natural products identified for example from soil-living actinomycetes or myxobacteria or plant-associated pseudomonads.

Our goal is to use entomopathogenic *Xenorhabdus* and *Photorhabdus* bacteria, which live in symbiosis with entomopathogenic *Steinernema* and *Heterorhabditis* nematodes, respectively,^[6–8]^ as models to study the true ecological function of their bacterial natural products, since all interacting organisms can be studied easily in the lab. During these efforts, we have already identified chemically diverse compounds with various functions namely antibiotic, antifungal, or nematicidal activity or acting as signals between bacteria or even between bacteria and their nematode hosts.^[9]^ While we have focussed in the past on conserved natural products occurring in several or even all *Xenorhabdus* and *Photorhabdus* species,^[10,11]^ we have started to look in more detail into rare or even unique natural products or the respective BGCs trying to understand also their ecological function. A starting point for this work are BGCs that indeed encode resistance genes, suggesting antibiotic activity.

In bacteria, translation initiates with a formylated methionine, whose removal by peptide deformylases (PDFs), a metalloprotein essential for N-terminal methionine excision process, is critical for protein maturation.^[12,13]^ Due to their essential role and broad conservation, PDFs have emerged as promising antibacterial targets with actinonin being the first identified PDF inhibitor (PDI),^[14]^ prompting the development of various analogs^[15,16]^ and the identification of other natural PDIs.^[17,18]^

Here we describe the structure of a novel peptide/polyketide hybrid named xildivaline from *Xenorhabdus hominickii* acting as novel peptide deformylase inhibitor. Its biosynthesis includes the first described *trans*-methyltransferase in non-ribosomal peptide synthetases.

## Results and discussions

In our search for novel NPs with biological activity we identified a BGC in *Xenorhabdus hominickii* encoding a hybrid of non-ribosomal peptide synthetase (NRPS)^[19]^ and polyketide synthase (PKS)^[20]^ as well as a peptide deformylase XisD and a methyltransferase XisE (Fig. 1a, Table S1),^[21]^ which was found in 70 different Gammaproteobacteria strains highly conserved in *Vibrio* as evident from a CompArative GEne Cluster Analysis Toolbox (CAGECAT) analysis^[22]^ (Fig. 1b, Figure S1). The BGC is encoded on a large plasmid suggesting possible uptake or distribution between different bacteria.

**Figure 1.**
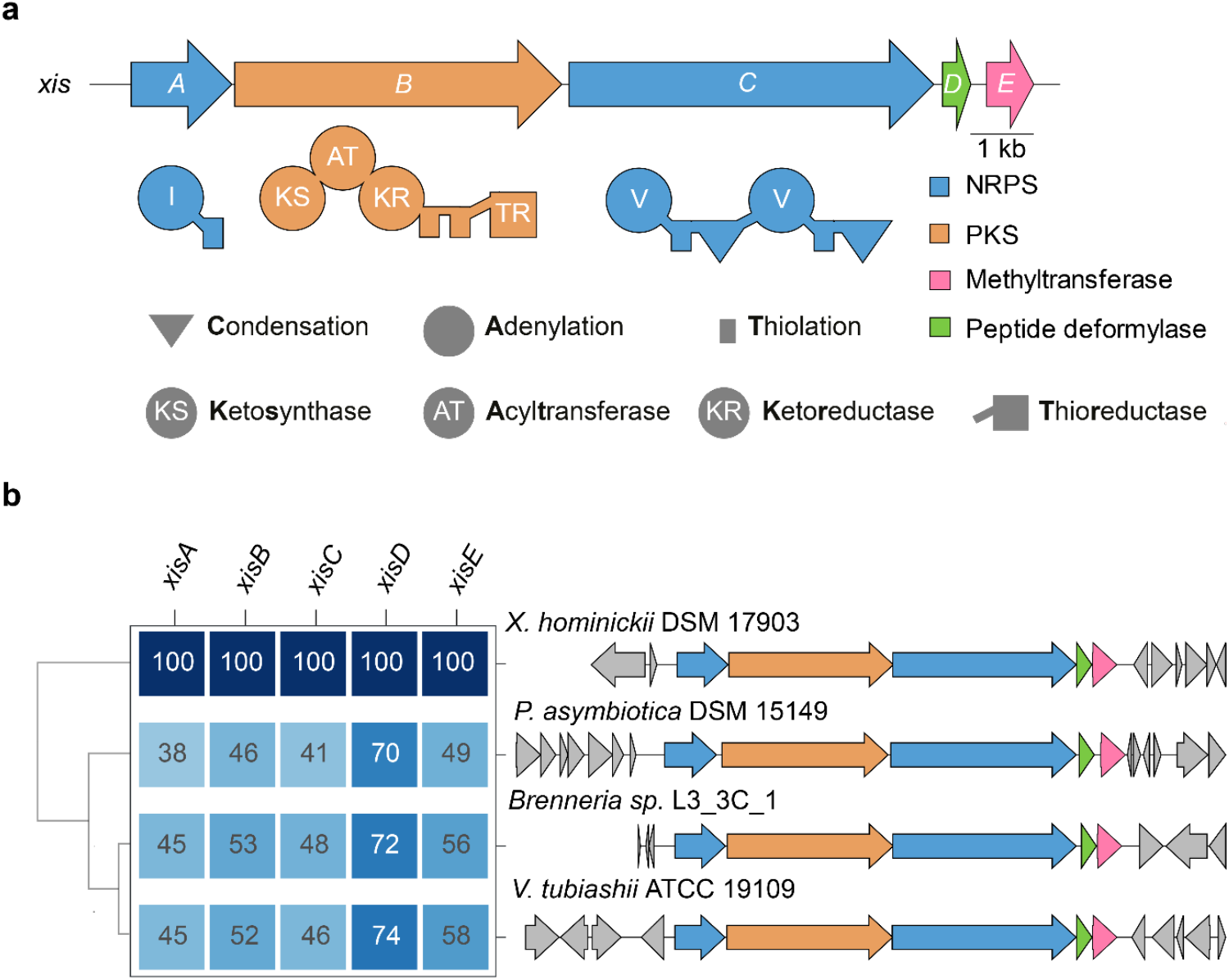
Xildivaline biosynthesis gene cluster (*xisABCDE*) from *X. hominickii* with domain organisation for NRPS and PKS enzymes (**a**) and similarity to BGCs from other bacteria (**b**).

Activation of the *xis* BGC in *X. hominickii* using a well-established promoter exchange strategy^[23,24]^ both in the wildtype (WT) or a natural product-reduced Δ*hfq* variant led to the production of several compounds hardly produced in the WT (Fig. 2a). Similarily, the complete BGC was also cloned on a pSEVA261b plasmid and expressed in *E. coli mtaA* (encoding the broad-specificity phosphopantetheinyl transferase MtaA^[25]^) leading to similar signals but at a much lower level (Fig. 2b). Additionally, we also expressed the almost identical BGC from *Vibrio tubiashii* DSM 19142 in *E. coli* also showing an almost identical production profile (Fig. S2).

**Figure 2.**
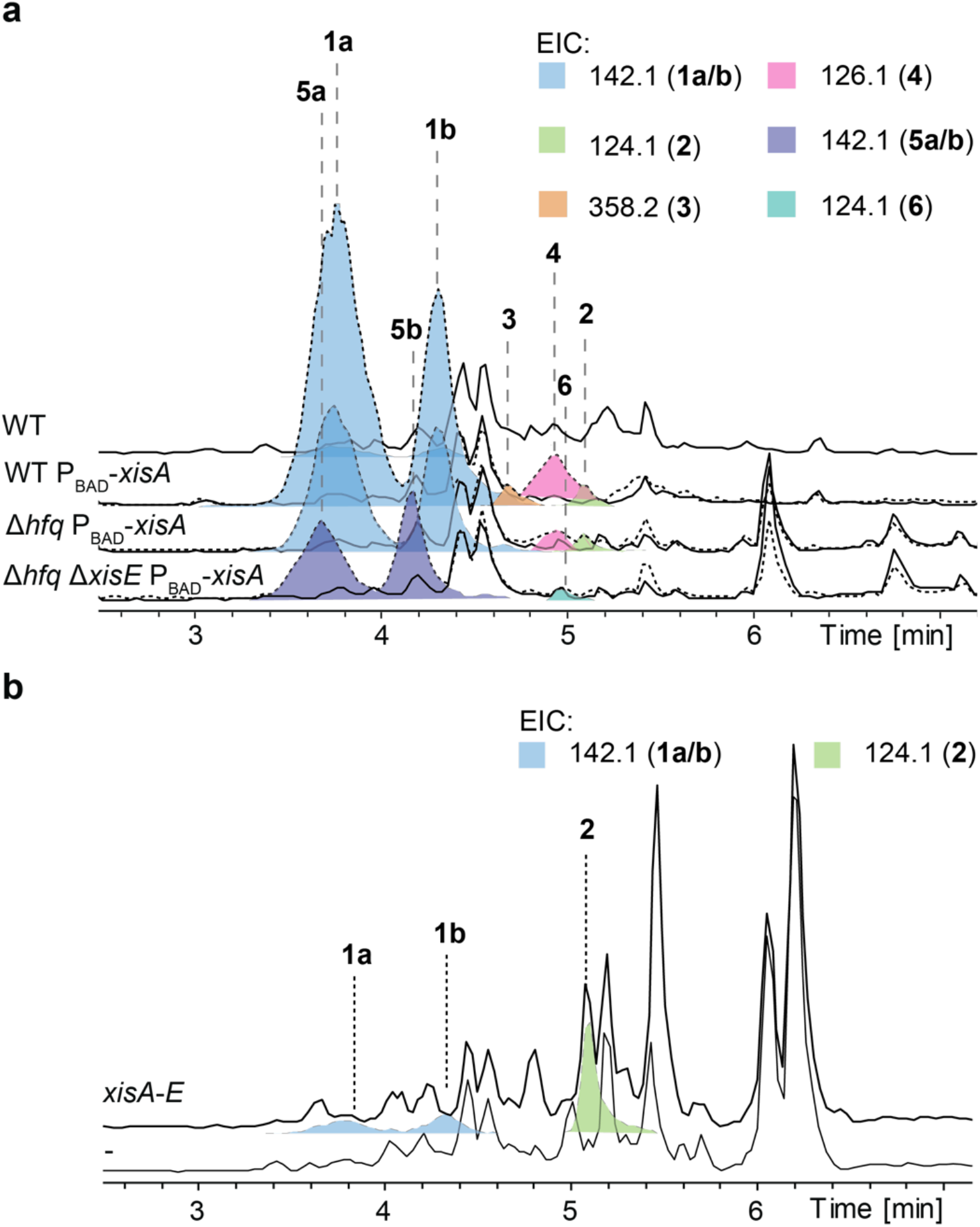
HPLC/MS chromatograms showing production of xildivaline derivatives in *X. hominickii* strains (**a**) and after heterologous expression of *xisABCDE* in *E. coli* (**b**). To highlight the EICs, the C-terminal fragment ions (see Figure S6) of the corresponding derivatives were selected, except for **3**.

The two main derivatives at all conditions **1a**/**1b** show a *m/z* 372.2856 suggesting the sum formula C_19_H_38_N_3_O_4_ [M+H]^+^ (Table S2) and MS fragmentation analysis indicate it to be a tripeptide of Val-Val-Ile extended by one polyketide unit and carrying one S-adenosyl methionine derived methyl group (Fig. 3, Fig. S3), which was confirmed by labeling experiments with *methyl*-[D_3_]-Met, [^12^C_5_]-Val or [^12^C_6_]-Ile fed to a fully ^13^C-labelled culture (Fig. S4) and a mass shift of 3 Da from differential cultivation in fully labeled ^14^N and ^15^N medium (Fig. S5).^[26]^ The MS-fragmentation of the D_3_-labelled **1a** suggests the methyl group to be located at the N-terminus (Fig. S6) as it was also suggested from the Val-Ile-polyketide fragment showing the non-methylated mass (Fig. S3 & S6). Therefore, XisD might be a member of the rare cases^[27,28]^ where the methyl group is not incorporated by the methyltransferase embedded in the NRPS assembly line^[19]^ but by acting in *trans*.

**Figure 3.**
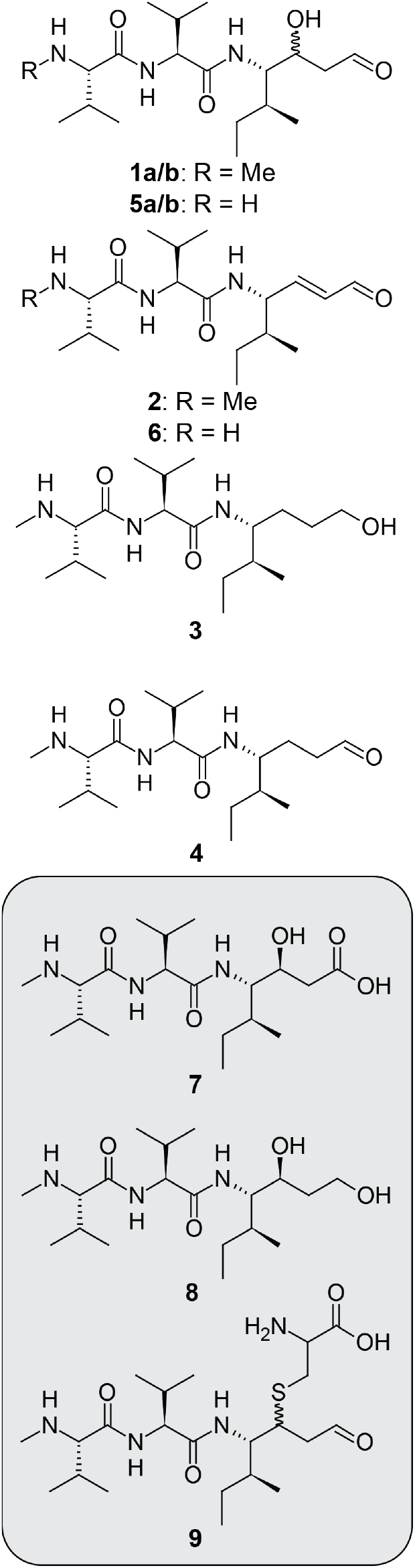
Proposed structures of identified xildivaline derivatives. **7**-**9** represent minor derivatives (see Fig. S7).

The C-terminus was identified to contain an aldehyde as it was predicted already from the thioester reductase (TR) domain (Fig. 1a).

While all other derivatives share the same peptide N-terminal part, they differ in the C-terminus, which might differ in the configuration of the hydroxy group (**1b**), or show a double bond instead of the hydroxy group (**2**) or are reduced showing a C-terminal alcohol (**3**) or an aldehyde (**4**) (Fig. 3; Table S2). Interestingly, **3** is neither found in the Δ*hfq* mutant nor the *E. coli* expression host suggesting that the additional reductase(s) responsible for aldehyde and double bond reduction might only be active in the WT (Fig. 2). A detailed analysis of a concentrated WT P_BAD_-*xisA* extract (Fig. S7) revealed minor amounts of additional derivatives; one carrying a C-terminal carboxylic acid group (**7**), probably due to premature release from one of the terminal T domains prior to thioester reduction, a derivative of **3** with a hydroxy group (**8**), as well as a cysteine adduct at the Michael system (**9**) (Fig. S7, Table S1). The structure of **7** was further confirmed by expressing the *xis* BGC with an engineered^[29]^ XisB containing a T-TE from the xenotetrapeptide producing XtpS NRPS^[30]^ instead of the original T-TR, indeed leading to the production of **7** instead of **1a/b** (Fig. S8).

For prediction of the hydroxy group configuration derived from the PKS ketoreductase (KR) domain,^[31]^ its amino acid sequence was compared to other KR domains suggesting it to be *S*-specific (Fig. S9). However, the fact that **1** and **9** were found as two peaks showing the same *m/z* suggests that they might be derived from Michael addition of water or cysteine to the double bond in **2**, which might result from **1** by spontaneous elimination of water.

Indeed, all attempts to isolate **1** or **2** failed, probably because of its rapid elimination of water generating an unsaturated aldehyde, which can react readily with nucleophiles like water, cysteine or 4-bromothiophenole (BTP) (Fig. S10). Therefore, we decided to analyze the bioactivity of xildivaline directly in the producing *E. coli* strain revealing that both XisD and XisE are important for protection against xildivaline as a deletion of either of these genes results in reduced growth (Fig. 4a).

**Figure 4.**
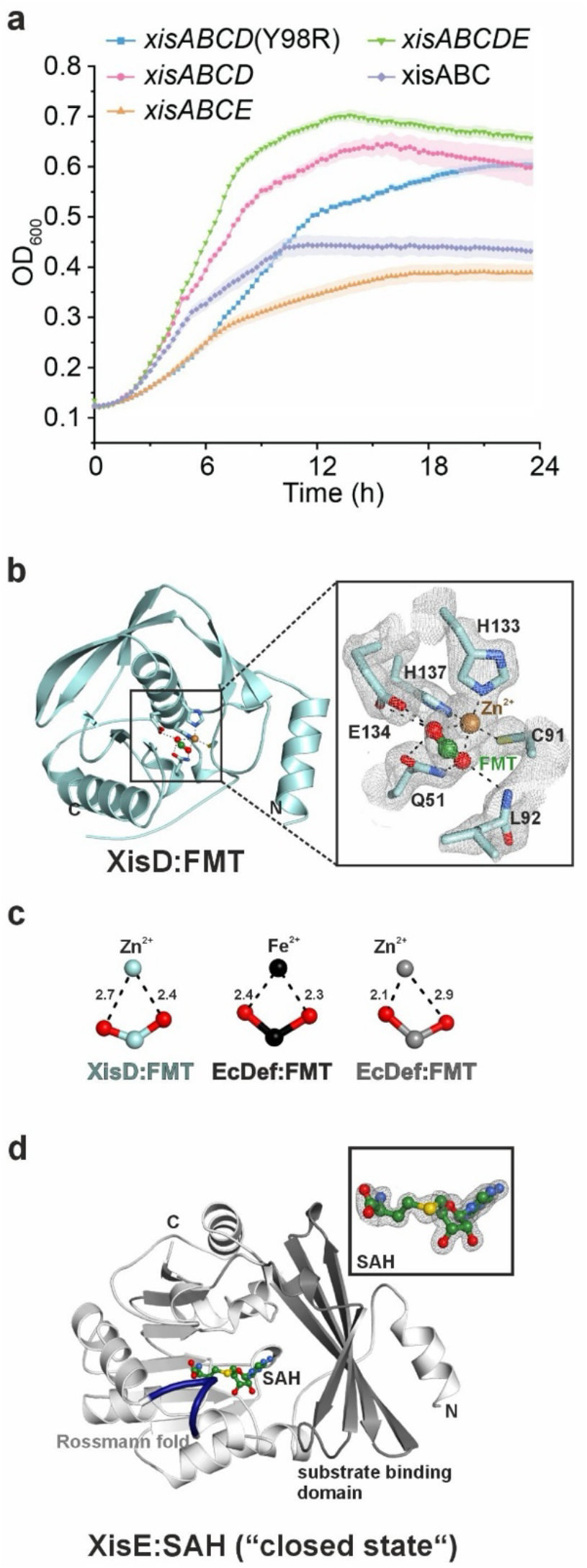
(**a**) Growth curves of *E. coli* strains expressing different combinations of *xis* genes. (b) Left: Ribbon illustration of XisD with bound formiate (FMT). N- and C-termini are labelled. Right: Zoom-in at the XisD active site. Key residues coordinating the Zn^2+^ ion (brown sphere) and the FMT (green and red ball-and-stick model) are shown as sticks with their 2F_O_-F_C_ electron density map (gray mesh contoured to 1σ; with Zn^2+^, FMT, amino acid side chains as well as main and side chain atoms of Leu92 omitted for phasing). Hydrogen bonds are indicated by black dotted lines. Distances are given in Fig. S12c. (**c**) Comparison of FMT binding to XisD:Zn^2+^ (light blue) as well as to EcDef:Fe^2+^ (black; PDB entry 1XEN (Jain *et al*., 2005)) and EcDef:Zn^2+^ (gray; PDB entry 1XEM (Jain *et al*., 2005)). Distances are indicated by black dotted lines and given in Å. (**d**) Ribbon structure of XisE in the “closed state”. Residues 38-44 (thickened purple segment) are well defined and form a substrate binding loop that shapes the active site pocket. The 2F_O_-F_C_ omit electron density map for the cofactor SAH (gray mesh contoured to 1σ) is shown in the right upper corner. See also Fig. S15.

Subsequently, we solved the structure of both proteins by X-ray crystallography (Table S4; for details see materials and methods as well as supplementary notes). XisD adopts the characteristic fold of class I peptide deformylases (Table S5). It has a deep and negatively charged active site pocket at the bottom of which a catalytic Zn^2+^ ion is chelated by residues Cys91, His133, His137 and a formiate molecule (Fig. 4b, Fig. S11a, Fig. S12a-d). Strikingly formiate binds the zinc ion in a bidentate manner (Fig. 4c, Fig. S12b-c, e). Zn^2+^ has previously been associated with a monodentate coordination by formiate and a lower catalytic activity compared to Fe^2+^ loaded *E. coli* deformylase.^[15]^ We therefore propose that XisD, although using a catalytic Zn^2+^ ion, coordinates formiate in a bidentate manner and might therefore be as active as iron containing peptide deformylases. Homologs of XisD are found in several Gammaproteobacteria encoding xildivaline-like BGCs (Fig. S13) and its deformylase activity can indeed substitute for canonical PDFs (Fig. S14).

The structure of XisE is typical of a SAM dependent class I methyltransferase. Between the Rossmann fold and the substrate binding domain there is a pronounced active site cleft which can be partly covered by a flexible loop (residues 38-44) (Fig. 4d, Fig. S15). The pocket lacks a basic residue for substrate deprotonation, but Phe133 could impact catalysis by stabilizing the intermediate positive charge at the transferred methyl group by its aromatic system. XisE is structurally related to celesticetin methyltransferase CcbJ^[32]^ and to the cypemycin N-terminal methyltransferase CypM^[33]^ (Table S6), but the catalytic residues of CcbJ (Tyr9, Tyr17 and Phe117)^[32]^ are not conserved in XisE (Fig. S11b). Moreover, the key secondary structure element of the substrate binding lobe, a four-stranded antiparallel β-sheet, is more extended in XisE, thereby increasing the active site cleft in comparison to CcbJ and CypM (Fig. S16a-c) and suggesting a larger substrate. While CcbJ and CypM oligomerize via the 4-stranded antiparallel β-sheet (Fig. S16b-c), XisE-related monomeric methyltransferases (for instance PDB entries 1Y8C and 3D2L, Table S6, Fig. S16d-e) share the inwards orientation of the β-strands, suggesting a role in shaping the substrate binding pocket rather than oligomerization. Since no non-methylated xildivaline derivatives were produced in the WT strain (Fig. 2 & S7) and because of the relatively large active site cleft XisE might act on the XisA-, XisB- or XisB-bound intermediates as proposed in Fig. 5 and not on free derivatives **5a/b**.

**Figure 5.**
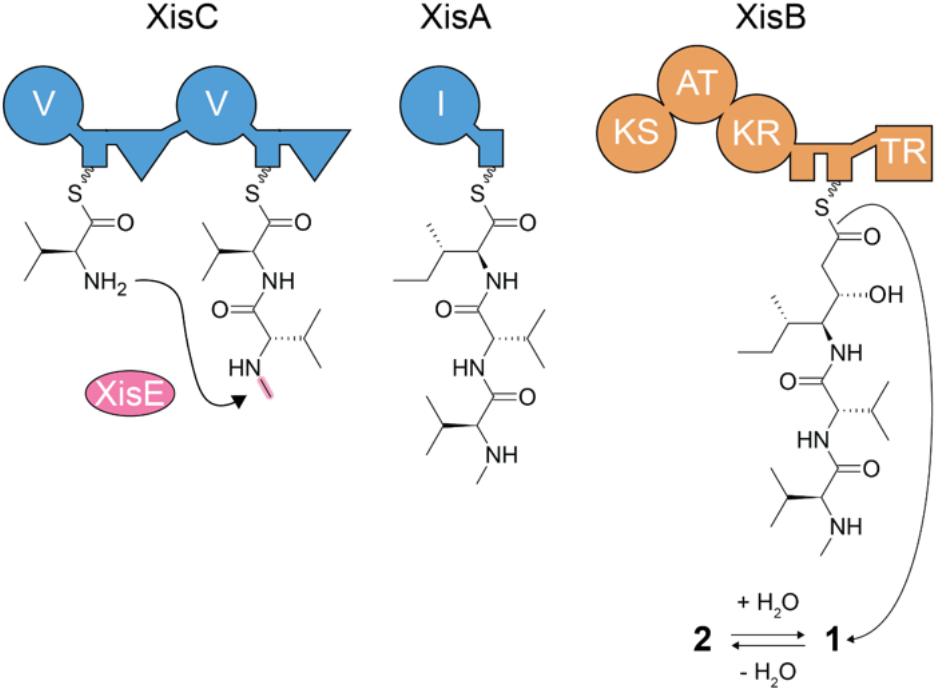
Proposed biosynthesis of xildivaline.

To investigate the interaction between xildivaline and its putative molecular target, we used AlphaFold (v3.0.0)^[34]^ to model the structure of bacterial peptide deformylases (PDFs) in complex with Fe^2+^ and xildivaline analogs (**1a, 2, 4a**, and **5**). Alphafold prediction of XisD was very similar to the crystallography data (Fig. 4b; 0,35 Å RMSD over 139/160 carbons present in the crystal structure). All ligands were consistently predicted to bind within the catalytic pocket, with their aldehyde moiety positioned near the divalent metal ion and key catalytic residues, Cys91 (motif II) and His133/His137 (motif III), which coordinate the metal and activate the nucleophilic water molecule (Fig. S11). This conformation mirrors the positioning of the natural formylated methionine during catalysis,^[35,36]^ supporting a substrate-mimetic mode of binding for xildivaline. Predicted hydrogen bonding patterns revealed both conserved and divergent features between *E. coli* PDF and XisD. While several residues, such as Ile45, Gly46, Gln51, and Gly90, were involved in ligand binding in both enzymes, suggesting a shared core interaction network, key differences were also observed.

In *E. coli* PDF, Arg98 was predicted to form a hydrogen bond with the amine group of the non-methylated derivative **4a**. This interaction was not detected with other xildivaline analogs, likely because the arginine side chain sits slightly beyond hydrogen bond distance (4.0–4.2 Å). Still, functional data indicate that Arg98 influences ligand binding, as its presence impairs *E. coli* growth upon BGC induction, suggesting a role in sensitivity to xildivaline (Fig. 4a). In contrast, XisD lacks Arg98 and instead encodes a tyrosine at the same position, which is predicted to point away from the ligand. This is consistent with structural data from the Def2 PDF of *V. cholerae* bound to actinonin.^[37]^ Additionally, XisD contains a unique Tyr126 (replaced by Leu in *E. coli* PDF) that forms a predicted hydrogen bond with the ligand. Another notable residue is Glu88, conserved in both enzymes. While it was predicted to interact with the ligand’s amine in XisD only, such an interaction could plausibly occur in *E. coli* PDF as well, especially at physiological pH, where the protonated amine may form an ionic bond not fully captured by static modeling.

## Conclusion

We could confirm the structure and biosynthesis of a new type of PDF inhibitor, which suggests a *trans*-methylation during the NRPS/PKS-based biosynthesis resulting in highly reactive compound named xildivaline (**1**). While α-N-methylation in *trans* has been described for ribosomally-derived peptides^[33,38]^ and proteins in bacteria^[39]^ and eukarya^[40–42]^, we are not aware of any example for NRPS-derived compounds, where α-N-methylation usually is derived from methyltransferases embedded in the NRPS as in the case of teixobactin.^[43]^ **1** rapidly eliminates water to form probably an even more active PDF inibitor **2** which can be reduced to **3** by alcohol dehydrogenases not yet identified. During our final experiments to solve the biosynthesis of xildivaline, the structurally highly similar gammanonin from heterologous expression of the *V. tubiashii* BGC in *E. coli* was described.^[44]^ It is a leucine instead of isoleucine variant of **7** showing no PDF activity probably due to the reduction of the aldehyde warhead. However, its OH configuration was determined by NMR as *S* confirming *S* configuration in xildivaline for the first product released from the enzyme machinery. Since the BGCs from *X. hominickii* and *V. tubiashii* are indeed very similar, we tested whether also xildivaline derivatives containing Leu instead of Ile are produced and indeed we could identify such derivatives albeit at much lower amounts (Fig. S17).

Despite the instability of xildivaline, our work might pave the way to develop novel PDF inhibitors not dependent on the hydroxamate moiety as in actinonin, which is highly reactive but also quite toxic *in vivo*. Here, an α,β-unsaturated carboxylic acid or methyl ester might be an interesting derivative for further studies, which can probably even obtained by NRPS engineering as shown during the engineering of novel syrbactin derivatives.^[45]^

It was recently shown that many bacterial species encode multiple peptide deformylase (PDF) genes, and that these accessory PDFs can contribute to resistance against natural PDF inhibitors found in the environment, potentially influencing microbial competition in ecological niches.^[37]^ For instance, *Vibrio cholerae* harbors an accessory PDF that confers strong resistance to actinonin, produced by *Streptomyces* species.^[37]^ However, given that *Streptomyces* and *Vibrio* typically occupy distinct ecological habitats, it seems unlikely that actinonin itself was the selective pressure that drove the acquisition or maintenance of this resistance mechanism in *Vibrio*.

In contrast, the wide distribution of the xildivaline BGC among *Vibrio* species suggest that the presence of these compounds in the environment may have exerted a more direct and ecologically relevant selective pressure. Indeed, it has been shown that a xildivaline-like natural product produced by *Vibrio ordalii* 12-B09 is capable of antagonizing >25% of other *Vibrio* isolates^[46]^ and recently it has been shown that it additionally can trigger prophage induction and subsequent lysis of prophage-encoding *Vibrio* strains in sublethal concentrations,^[47]^ suggesting an additional competitive advantage and thus a new natural product function beyond simple antibiotic activity.

## Author contributions

A.R., M.W. and J.C. constructed all strains and performed all experiments, except E. coli expressing the *V. tubiashii* BGC generated by C. J.. Crystallization and structure elucidation of XisD and XisE were performed by E.H. and M.G. and analysis of XisD, comparison with other PDFs and xildivaline/PDF modelling were performed by M.L. and D.M.. A.R. and H.B.B. wrote the paper with input from all authors.

## Acknowledgements

Work in the Bode lab was supported by an ERC Advanced Grant (835108) and the Max Planck Society. Work in the Mazel lab was supported by the Fondation pour la Recherche Médicale Equipe FRM (EQU202103012569 and FDM202106013531, to M. Lambérioux), the Institut Pasteur and the Centre National de la Recherche Scientifique. The synchrotron data was collected at beamline P13 operated by EMBL Hamburg at the PETRA III storage ring (DESY, Hamburg, Germany) under grant no. MX-970. We thank the staff of beamline P13 for assistance during data collection and the student A. Tandinata for experimental support. M.L. and D.M. would like to thank the Institut Pasteur strain collection (CIP) for providing *V. tubiashii* strains and Magaly Ducos-Galand for technical assistance.

## Data availability statement

All data and materials can be found within the manuscript, supporting information or can be requested from the corresponding authors.

