## Supplementary methods, Tables and Figures for "Identification and biosynthesis of xildivaline, a novel and widespread peptide deformylase inhibitor from Gammaproteobacteria"

### Supplementary information

### A. Materials & Methods

#### Cultivation of strains

Microorganisms were cultivated in LB medium for cloning purposes and overnight cultures. For cultivation on LB agar plates, 1.5 % w/v) agar was added. If not stated otherwise, *E. coli* cells were grown at 37 °C while *Xenorhabdus* was incubated at 28 °C. Antibiotics kanamycin (50 µg/mL for *E. coli* and 25 µg/ml for *Xenorhabdus*) or gentamycin (20 µg/mL for *E. coli* or 10 µg/ml for *Xenorhabdus*) were added when appropriate. Production cultures were cultivated in XPP medium [1] at 28 °C for 72 h. For induction, 0.2 % (w/v) L-arabinose was added at the beginning of cultivation.
Vibrio cholerae strains were grown in Mueller-Hinton (MH) broth. When required, antibiotics were used at the following final concentrations: carbenicillin (100 µg/mL) and kanamycin (25 µg/mL). The *E. coli* donor strain NGEpir was cultured in MH broth supplemented with 0.3 mM diaminopimelic acid (DAP).

#### Promoter exchange and deletions in *X. hominickii*

Promotor exchange was performed according to the described method in [1]. The promoter exchange mutant *X. hominickii* $P_{BAD}$_*xisA* was generated via conjugation between *X. hominickii* WT or its Δ*hfq* variant and donor strain *E. coli* ST18 + pCEP_kan_xisA. Gene deletions were performed according to the method described in [2]. Approximately 1 kb long regions flanking the target gene were amplified via PCR and thereby overhangs to the other flanking fragment and the target plasmid pEB17 were introduced. The confirmed plasmid was transformed into *E. coli* ST18 and this strain was then used as donor strain for conjugation with the recipient *Xenorhabdus* strain. 5 mL LB cultures were inoculated from overnight cultures in two different ratios: 1:50 and 1:100 for *E. coli* and 1:25 and 1:50 for *Xenorhabdus*, respectively. After growth to an OD595= 0.6, 2 mL of culture were harvested for each *E. coli* and *Xenorhabdus* by centrifugation. The harvested cells were washed twice with 1 mL fresh LB at 8000 rpm for 1 min and resuspended in 200 µL LB. Donor and recipient strain were mixed in a droplet on an LB agar plate and incubated overnight at 28 °C. The next day, the cell mass was scraped from the LB agar plate, resuspended in 2 mL LB and plated on two large LB agar plates containing kanamycin (50 µg/mL). Following incubation at 28 °C for two days, overnight cultures with and without antibiotics were prepared for grown colonies. From cultures grown in plain LB medium, serial dilutions were plated on LB agar plates containing 6 % w/v sucrose. Successful deletions were verified by PCR.

#### Growth experiments

Growth phase monitoring was conducted in 96-well microtiterplates using a Spark® microplate reader (TECAN) with orbital shaking at 22 °C for 24 h. OD595 was measured every 15 minutes. LB medium (with appropriate antibiotics) was used for cultivation and culture wells were inoculated at OD595 = 0.1 from overnight culture. For induction, 0.2 % (w/v) L-arabinose was added at the beginning of cultivation.

**Bacterial Conjugation**

Plasmid transfer into *V. cholerae* was performed by conjugation from *E. coli* NGEpir, which carries the RK2 conjugation machinery integrated into its chromosome, as described previously [3].

**Phylogenetic Analysis of Peptide Deformylases**

Amino acid sequences of peptide deformylases (PDFs) were aligned using MAFFT (v7.467, option --auto). [4] Phylogenetic reconstruction was carried out with IQ-TREE (v2.2.2.2) using the -m TEST function to select the best-fitting model according to the Bayesian Information Criterion (BIC) [5]. Branch support was assessed with 1,000 ultrafast bootstrap replicates. [6]

**In Vivo Assay of PDF Activity in *Vibrio cholerae***

The enzymatic activity of heterologous PDFs was assessed in a *V. cholerae* Δ*def1* Δ*def2* mutant, in which essential deformylase function was maintained by a constitutively expressed def2 allele on a temperature-sensitive low-copy plasmid. A second plasmid encoding the PDF of interest was introduced. Growth at 42 °C served as a readout for in vivo deformylase activity, as only strains expressing an active PDF from the second plasmid can proliferate at the restrictive temperature. For more details see: [3].

**Cloning**

Genomic DNA was purified using either the Gentra Puregene Yeast/Bact. Kit (Qiagen), or the Monarch® Genomic DNA Purification Kit (NEB), according to the manufacturers' protocols. Polymerase chain reaction were performed with Q5 High-Fidelity DNA Polymerase according to the manufacturers' protocols.

Plasmid with an oriV(SC101) origin of replication were derived from pMP394, with pBAD promoter and araC removed. Plasmids with an oriV(RK2) origin of replication were derived from pSEVA228 [7], with xylS removed. Linearized pSEVA261b or pSEVA631b expression vectores were obtained by PCR. Inserts of the *xis* BGC were amplified with the listed primers in table *X* from *X. homickii* genomic DNA or already assembled vectors. TR-TE exchange were performed according to [8] by using XUT site I.

Plasmid assemblies were performed using Gibson Assembly as described in [9] and [10] or with NEBuilder® HiFi DNA Assembly Master Mix according to the manufacturers' protocols.

**LC-MS analysis**

After three days of cultivation of xildivaline producing strains the cultures were harvested by mixing with methanol in a ratio of 1:100 and processed as described [11]. For 4-bromothiophenol (BTP) addition, 50 µl (1 mg/mL stock in MeOH) 4-bromothiophenol were added to 500 µL thawed culture supernatant. Samples were incubated overnight at 30 °C. As controls, samples without BTP addition and samples with addition of 500 µL MeOH were used. Prior to HPLC-MS analysis, samples were diluted 1:10 with ACN and centrifuged at full speed for 30 min.

**Heterologous expression of XisD and XisE**

The coding sequences of XisD and XisE were ordered as codon optimized synthetic genes (Table S3) and cloned with restriction enzymes *Bam*HI-HF and *Pst*I-HF in a pET-Duet-1 based vector providing a N-terminal $His_6$-SUMO tag (MWG Eurofins Genomics, Ebersberg). After transformation of the resulting plasmids in *Escherichia coli* BL21-Gold (DE3) cells, a preculture in LB medium supplemented with 180 mg/L ampicillin was inoculated with a single transformant and incubated overnight at 37 °C, 130 rpm. The next day, the main culture (2-6 L of LB medium supplemented with 180 mg/L ampicillin) was started by addition of preculture in a 1:50 ratio. The main culture was incubated at 37°C, 130 rpm until an optical density ($OD_{600nm}$) of 0.5-0.7 was reached. The flasks were then cooled to 20 °C and 0.5 mM isopropyl-β-D-1-thiogalactopyranoside was added to induce gene expression. After 16-20 h at 20 °C and 130 rpm, cells were harvested by centrifugation, washed with 0.9% (w/v) NaCl and frozen at -20 °C until further use.

**Purification of recombinantly produced XisD and XisE**

Cells were resuspended in buffer A (100 mM Tris/HCl pH 7.5, 500 mM NaCl, 20 mM imidazole, and 2 mM β-mercaptoethanol) and lysed by sonication. After centrifugation at 41,000 x g for 30 min at 4 °C, the supernatant was loaded on a 5 mL nickel chelating Sepharose HP column (Cytiva), pre-equilibrated with buffer A (flow rate 5 mL/min). After washing with buffer A, the protein was eluted by applying a gradient 0% to 100% buffer B (100 mM Tris/HCl pH 7.5, 500 mM NaCl, 500 mM imidazole, and 2 mM β-mercaptoethanol) over 10 column volumes (CV). Fractions containing the protein of interest were pooled and supplemented with $His_6$-tagged SUMO-protease to remove the tag from the target protein. The mixture of protein and protease was dialyzed overnight at 4 °C against buffer C (20 mM Tris/HCl, pH 7.5, 100 mM NaCl, and 2 mM β-mercaptoethanol). Next, the protein was again loaded on a 5 mL nickel chelating Sepharose HP column, pre-equilibrated with buffer A to remove the protease and the cleaved tag. The flow through, containing the protein of interest, was collected, concentrated with a 10 kDa cutoff filter, and further purified using size exclusion chromatography. To this end, a Superdex 200 16/60 column (GE Healthcare) was equilibrated with 1.2 CV of buffer D (20 mM Tris/HCl, pH 7.5, 100 mM NaCl, 2 mM dithiothreitol). The concentrated protein solution was applied via a capillary loop and eluted with an additional 1.2 CV of buffer D. Fractions were collected and probed for purity by SDS-PAGE.

**Crystallization and structure determination of XisD**

Purified XisD (50 mg/ml) was subjected to sitting drop vapor diffusion crystallization by mixing different ratios of protein with reservoir solutions and incubating the crystal trays at 20 °C. Crystals grew from 0.01 M zinc sulfate, 0.1 M MES pH 6.5 and 25% (w/v) PEG550 MME and were cryoprotected by the addition of a 1:1 mixture of 50% (v/v) glycerol and reservoir before vitrification in liquid nitrogen.

Diffraction data were collected at the beamline P13, DESY, Hamburg, Germany. Reflection intensities were evaluated with XDS. [12] The X-ray structure of XisD was solved by molecular replacement calculations (Phaser program within CCP4 [13] with its AlphaFold3 model [14]. After refinements of the model with Coot (Emsley et al., 2010) and REFMAC5 [15] water molecules were placed with ARP/wARP solvent [16]. Final refinements with REFMAC5 resulted in reasonable R-values and geometry as confirmed by PROCHECK [17] and MolProbity [18].

**Crystallization and structure determination of XisE**

Purified XisE was concentrated to 33 mg/ml, supplemented with 2 mM SAM (dissolved in water) and used for sitting drop vapor diffusion crystallization trials at 20 °C. Crystals for the open state grew from a 2:1 ration of protein and reservoir containing 0.2 M magnesium sulfate, 20% (w/v) PEG4000, and 10% (v/v) glycerol, while the closed state of XisE was obtained from 0.2 M calcium acetate and 20% (w/v) PEG3350. Crystals were cryoprotected with a 1:1 mixture of 50% (v/v) glycerol and reservoir and vitrified in liquid nitrogen.

Diffraction data were recorded at DESY (beamline P13, Hamburg, Germany) and evaluated with XDS [12]. The structure of XisE was solved by molecular replacement with Phaser [19] (CCP4 suite) using the AlphaFold3 [14] prediction of XisE as a search model. The model was iteratively refined with Coot [20] and REFMAC5 [15] and water molecules were placed with ARP/wARP solvent [21]. TLS refinements finally yielded excellent R-values and geometry. The model was proven to fulfill the Ramachandran plot using PROCHECK [17] and evaluated by MolProbity [18].

### B. Supplementary Notes

#### Supplementary note on XisD

XisD was expressed as a N-terminal $His_6$-SUMO fusion in *Escherichia coli*. The protein was purified by affinity and size exclusion chromatography with tag removal in between and crystallized. The structure of XisD was solved to 1.65 Å resolution with two monomers per asymmetric unit (Table S4). The compact fold of XisD is typical of class I peptide deformylases (PDFs) from Gram-negative bacteria and a homology search revealed tight relationships to corresponding enzymes from *Vibrio cholerae* and *Escherichia coli* (Table S5). XisD also shares the typical sequence motifs of PDFs (GϕGϕAAXQ, EGCϕS, and HEϕDH with ϕ being a hydrophobic and X any amino acid; Fig. S11a) [22]. Like other PDFs XisD uses a catalytic metal coordinated by Cys91, His133 and His137 for catalysis (Fig. 4b, Fig. S12a-c). Notably, deformylases exhibit a pronounced metal promiscuity and enzymatic activity has been reported for zinc-, iron-, cobalt-, manganese- as well as nickel-loaded variants [22], leaving it enigmatic which metal ion is *in vivo* the natural cofactor. X-ray fluorescence analysis of various XisD crystals revealed the presence of zinc ions independent of the crystallization condition. Although formiate (FMT) was neither added to the protein preparation or during crystallization, we identified this ligand at the bottom of a pronounced negatively charged active site pocket of both XisD monomers (Fig. S12d). FMT coordinates the active site metal and hydrogen bonds to the side chains of Gln51 and Glu134 as well as to the Leu92NH (Fig. S12b-c). Previous studies proposed differences in formiate coordination between $Zn^{2+}$ (monodentate) and $Fe^{2+}$ (bidentate) loaded *E. coli* peptide deformylase to account for the lower activity of $Zn^{2+}$ containing enzyme [23]. Notably, we here report for zinc bound XisE a bidentate coordination of formiate, which is only slightly distorted and highly similar to iron-loaded *E. coli* peptide deformylase [23] (Fig. 4c and Fig. S12e). Since *E. coli* deformylase is highly stable in the presence of zinc [24] but less active due to the monodentate coordination of formiate [23], it is tempting to speculate that XisD might be as active as iron-containing deformylases but more stable and hence excellently fulfill its function as a resistance protein against xildivalines.

#### Supplementary note on XisE

Heterologous production and purification of XisE were carried out as for XisD. The purified protein was crystallized and the structure of XisE with its cofactor SAH was solved to 1.4 Å resolution (Table S4). A Dali search identified the celesticetin methyltransferase CcbJ [25] and cypemycin N-terminal methyltransferase CypM [26] as closest structural homologues of XisE (Table S6, Fig. S11b, Fig. S16). CcbJ and CypM share a similar subunit fold with XisE but are hexamers, while XisE is a monomer. The two-domain topology of XisE corresponds to that of a typical SAM dependent class I methyltransferase (Fig. 4d and Fig. S15). Between the Rossmann fold and the substrate lobe, there is a pronounced and solvent-accessible substrate binding cleft. As part of the Rossmann fold XisE encodes the characteristic GxGxG motif and an acidic residue at the end of β-sheet 2 (here Asp93) involved in cofactor binding (Fig. S11b). The active site cavity is spacious and solvent exposed. Although it appears negatively charged (Fig. S15), it is lined with numerous hydrophobic residues (Fig. S11b). Residues 38-44 form a flexible putative substrate binding loop that is either ordered (closed conformation, Fig. S15b) or disordered (open conformation, increased B-factors, Fig. S15a) depending on the crystallization condition. In the latter case also the homocysteine moiety of the SAH has increased B-factors (Fig. S15). Notably, the active site does not encode any obvious basic residue that could facilitate substrate deprotonation. Phe133 however occupies a prominent position that could impact catalysis, for instance by stabilizing the intermediate positive charge at the transferred methyl group via its aromatic system. Although XisE shows highest structural homology to CcbJ, the catalytic residues of CcbJ (Tyr9, Tyr17 and Phe117 [25]) are not conserved in XisE (Fig. S11b). Comparison of the 3D structures of CcbJ, CypM and XisE reveals that the Rossmann domain is conserved but that there are significant differences in the substrate binding domain (Fig. S16b-d). While the key secondary structure element of the substrate binding domain, a four-stranded antiparallel β-sheet, is present in CcbJ, CypM and XisE, their sequence (Fig. S11b) and position relative to the Rossmain domain is different (Fig. S16a-c). In CcbJ and CypM the β-sheet is turned towards the Rossmann domain, while in XisE it is extended, thereby increasing the active site cleft. Since CcbJ, CypM and XisE structures have been determined in the absence of ligand, the different orientation of the substrate binding domain is not a result of ligand recognition but rather an inherent structural feature of these methyltransferases. Strikingly, CcbJ and CypM oligomerize via the 4-stranded antiparallel β-sheet (Fig. S16b-c), but XisE-related monomeric methyltransferases (for instance PDB entry codes 1Y8C and 3D2L, Table S6, Fig. S16d-e) show the same inwards orientation of the β-strands, suggesting a role in shaping the substrate binding pocket rather than oligomerization.

### C. Supplementary Tables

#### Table S1.

| Gene | NCBI locus tag (NZ_NJAI01000013.1) | Size (bp) | Proposed coding sequence function |
|---|---|---|---|
| *xisA* | Xhom_RS23680 | 1704 | non-ribosomal peptide synthetase |
| *xisB* | Xhom_RS23685 | 5505 | type I polyketide synthase |
| *xisC* | Xhom_RS23690 | 6144 | non-ribosomal peptide synthetase |
| *xisD* | Xhom_RS23695 | 507 | peptide deformylase |
| *xisE* | Xhom_RS23700 | 819 | SAM-dependent methyltransferase |

**Table S2.** MS data of all identified xildivaline derivatives.

| Compound | Chemical formula | Detected $[M+H]^+$ | Calculated $[M+H]^+$ | Δppm | Retention time $t_R$ [min] |
|---|---|---|---|---|---|
| **1a** | $C_{19}H_{37}N_3O_4$ | 372.2858 | 372.2857 | 1.2 | 3.8 |
| **1b** | $C_{19}H_{37}N_3O_4$ | 372.2857 | 372.2857 | 1.5 | 4.3 |
| **2** | $C_{19}H_{35}N_3O_3$ | 354.2751 | 354.2751 | 0.0 | 5.1 |
| **3** | $C_{19}H_{39}N_3O_3$ | 358.3064 | 358.3064 | 0.5 | 5.0 |
| **4** | $C_{19}H_{36}N_3O_3$ | 356.2906 | 356.2908 | 0.2 | 4.9 |
| **5a** | $C_{18}H_{35}N_3O_4$ | 358.2700 | 358.2700 | 0.0 | 3.7 |
| **5b** | $C_{18}H_{35}N_3O_4$ | 358.2700 | 358.2700 | 0.0 | 4.2 |
| **6** | $C_{18}H_{33}N_3O_3$ | 340.2591 | 340.2595 | 1.0 | 5.0 |
| **7** | $C_{19}H_{37}N_3O_5$ | 388.2806 | 388.2806 | 0.0 | 3.9 |
| **8** | $C_{19}H_{39}N_3O_4$ | 374.3012 | 374.3013 | -0.2 | 3.7 |
| **9** | $C_{22}H_{42}N_4O_5S$ | 475.2946 | 475.2949 | -0.6 | 3.7 |
| **2-BTP** | $C_{35}H_{40}BrN_3O_3S$ | 542.19 | 542.2046 | a* | 8.8 |

**Table S3. Synthetic gene fragments.**

| Gene | Sequence (codon-optimized) 5' → 3' |
|---|---|
| *xisD* | ACCGTTCGTAAAATCATCGAAATCCCGGACGAACGTCTGCGTGTTACCTACCAGAAAGTTGAATGCGTTTCTACCGTTCAGAC<br>CCTGATCGACGACATGCTGGACACCGTTTACTCTACCGACCACGGTATCGGTCTGGCTGCTCCGCAGATCGGTCGTACCGAAG<br>CTGTTGCTATCATCGACATCTCTACCACCCGTGACAACCCGCTGATCCTGATCAACCCGGAACTGGTTGAAACCGACGGTGAA<br>TACATCGGTGAAGAAGGTTGCCTGTCTGTTCCGGGTTTCTACGCTAACGTTAAACGTTTCAAAAAAATCAAAGTTAAAGCTCT<br>GAACCGTGAAGGTGAAGAATTCTTCGTTGAAGACGACGGTTACCTGGCTATCGTTATGCAGCACGAAATCGACCACCTGCACG<br>GTAAAATCTTCATCGACTACCTGTCTCCGCTGAAACGTCAGATGGCTATGAAAAAAATCAAAAAACAGAAAATGATCAACAAC<br>AAATAA |
| *xisE* | AACACCGAAATCCTGAAAGACTTCCTGCCGGCTATCCGTTCTTCTGACTACATCATGGACTTCGGTGACCGTGCTTTCTCTCA<br>GCGTATGCTGAAAGAACACCTGAACCAGGGTTCTGAATTCGCTTCTCGTACCATCTCTGAAATCGACCGTCAGGTTTCTTTCC<br>TGTTCGACAAATACCTGACCCAGGGTGACAAACTGCTGGACCTGGGTTGCGGTCCGGGTCTGTACACTACTAGGTTCGCTGAA<br>AAAGGCGTTACCACCCTCGGCGTTGACGTAAGCCCGGCTGCTATCGAATACGCTAAAGAACACGCTACCTCTGCTGAAACCTA<br>CCAGCAGATCGACCTGGACAAATTCGACTCTAACGAACAGTTCGACCTGGTTCTGCTGCTGTTCGGTATCGCTAACAACCTGG<br>AACGTCTGGACACCCTGCTGCGTAAACTGAAACGTAACCTGAAATCTGGTGCTAAACTGGTTTTCGAACTGATGGACCTGGAA<br>TTCATGAAATCTCTGGAACAGGGTAATGGCACCTGGGTGTTCCACCCGGAAGGTGGTCTGCTGTCTGAACAGCCGCACTACCA<br>GCTGTGCCGTCGTGTTTGGTTCGAAGACCAGAAAACCCTGATCGACCGTAACATGGTTATCACCGACTCTGCTCAGACCTCTA<br>TGTACGAAGGTGTTTTTTTCGGCTTCGAACTGTACGACTTCAACCAGCTGCTGCAAAAAGCTGGTTACAAAGAAGCTCACATC<br>ATCTGCCGTCAGCTGGAAAAAGGTGAACTGACCAAACACTTCTTCATGGTTGAAACCGAACTGGCTTAA |

**Table S4.** X-ray data collection and refinement statistics for XisD and XisE structures.

| | ***XisD:FMT*** | ***XisE:SAH closed state*** | ***XisE:SAH open state*** |
|---|---|---|---|
| **Crystal parameters** | | | |
| Space group | $P2_1$ | P1 | P1 |
| Cell constants | a= 66.9 Å<br>b= 36.8 Å<br>c= 74.4 Å<br>β= 90.04° | a= 47.5 Å<br>b= 47.6 Å<br>c= 85.4 Å<br>α= 80.0°<br>β= 80.0°<br>γ= 73.6° | a= 47.5 Å<br>b= 47.6 Å<br>c= 85.4 Å<br>α= 80.4°<br>β= 80.2°<br>γ= 73.4° |
| Molecules / AU[a] | 2 | 2 | 2 |
| **Data collection** | | | |
| Beam line | P13, DESY | P13, DESY | P13, DESY |
| Wavelength (Å) | 0.976 | 0.976 | 0.976 |
| Resolution range (Å)[b] | 74-1.65<br>(1.75-1.65) | 45-1.4<br>(1.5-1.4) | 45-1.45<br>(1.55-1.45) |
| No. observations | 132938 | 525590 | 474279 |
| No. unique reflections[c] | 43076 | 132287 | 119221 |
| Completeness (%)[b] | 97.6 (98.0) | 96.0 (95.0) | 96.0 (95.3) |
| $R_{merge}$ (%)[b, d] | 6.8 (65.1) | 5.7 (56.9) | 4.5 (58.9) |
| I/σ (I)[b] | 9.0 (1.7) | 13.0 (2.9) | 15.2 (2.3) |
| **Refinement (REFMAC5)** | | | |
| Resolution range (Å) | 45-1.65 | 45-1.45 | 45-1.45 |
| No. refl. working set | 40919 | 125672 | 113260 |
| No. refl. test set | 2153 | 6615 | 5961 |
| No. non hydrogen | 2705 | 5148 | 4943 |
| Solvent | 114 | 540 | 434 |
| $R_{work}/R_{free}$ (%)[e] | 20.1/23.8 | 16.2/18.1 | 15.0/17.0 |
| r.m.s.d. bond (Å) / angle (°)[f] | 0.003/1.205 | 0.004/1.172 | 0.005/1.260 |
| Average B-factor ($Å^2$) | 31.1 | 20.1 | 24.1 |
| Ramachandran Plot (%)[g] | 97.2/2.8/0.0 | 98.1/1.9/0.0 | 98.7/1.3/0.0 |
| PDB entry code | 9RFP | 9RFR | 9RFS |

[a] Asymmetric unit

[b] The values in parentheses for resolution range, completeness, $R_{merge}$ and I/σ (I) correspond to the highest resolution shell

[c] Data reduction was carried out with XDS and from a single crystal
Friedel pairs were treated as identical reflections

[d] $R_{merge}(I) = \Sigma_{hkl}\Sigma_j \mid I(hkl)_j - \langle I(hkl)\rangle \mid / \Sigma_{hkl} \Sigma_j I(hkl)_j$, where $I(hkl)_j$ is the $j^{th}$ measurement of the intensity of reflection hkl and <I(hkl)> is the average intensity

[e] $R = \Sigma_{hkl} \mid |F_{obs}| - |F_{calc}| \mid / \Sigma_{hkl} |F_{obs}|$, where $R_{free}$ is calculated without a sigma cut off for a randomly chosen 5% of reflections, which were not used for structure refinement, and $R_{work}$ is calculated for the remaining reflections

[f] Deviations from ideal bond lengths/angles

[g] Percentage of residues in favored / allowed / outlier region

**Table S5.** Proteins structurally related to XisD according to Dali search.

| PDB entry code | Z-Score | R.m.s.d. [Å] | Identity [%] | Protein |
|---|---|---|---|---|
| 3qu1 | 28.6 | 0.9 | 60 | Peptide deformylase from *Vibrio cholerae* |
| 1g2a | 27.8 | 0.9 | 51 | The crystal structure of *E. coli* peptide deformylase complexed with actinonin |
| 5j46 | 26.8 | 1.0 | 43 | Crystal structure of a Peptide Deformylase from *Burkholderia multivorans* |
| 4wxl | 26.7 | 1.1 | 47 | Crystal structure of a peptide deformylase from *Haemophilus influenzae* complex with Actinonin |
| 3fwx | 26.7 | 1.2 | 50 | The crystal structure of the peptide deformylase from *Vibrio cholerae O1 biovar El Tor str. N16961* |
| 1n5n | 26.6 | 1.2 | 47 | Crystal Structure of Peptide Deformylase from *Pseudomonas aeruginosa* |
| 6caz | 26.3 | 1.2 | 51 | Crystal structure of a peptide deformylase from *Legionella pneumophila* |
| 3u04 | 26.2 | 1.3 | 38 | Crystal structure of peptide deformylase from *Ehrlichia chaffeensis* in complex with actinonin |
| 6jeu | 25.9 | 1.3 | 45 | K1U bound crystal peptide deformylase from *Acinetobacter baumanii* |
| 4e9b | 25.3 | 1.6 | 41 | Structure of Peptide Deformylase form *Helicobacter pylori* in complex with actinonin |

The Dali server [27] identified numerous structures with 3D similarity to XisD (search performed on the 16th of April 2025). The best 10 hits are shown and listed according to their Z-score. Redundant protein hits have been excluded from the list.

**Table S6.** Proteins structurally related to XisE (closed state) according to Dali search.

| PDB entry code | Z-Score | R.m.s.d. [Å] | Identity [%] | Protein |
|---|---|---|---|---|
| 4hh4 | 21.4 | 3.9 | 15 | Structure of the CcbJ Methyltransferase from *Streptomyces caelestis* |
| 7wzg | 19.9 | 3.6 | 17 | Cypemycin N-terminal methyltransferase CypM |
| 7zkh | 19.8 | 3.6 | 16 | C-Methyltransferase PsmD from *Streptomyces griseofuscus* with bound cofactor |
| 1y8c | 19.7 | 3.4 | 16 | Crystal structure of a S-adenosylmethionine-dependent methyltransferase from *Clostridium acetobutylicum* ATCC 824 |
| 6p3n | 19.6 | 4.2 | 15 | Tetrahydroprotoberberine N-methyltransferase in complex with S-adenosylmethionine |
| 6m81 | 19.3 | 34 | 16 | Crystal structure of TylM1 Y14F bound to SAH and dTDP-phenol |
| 5f8f | 19.2 | 2.9 | 16 | Crystal structure of Rv2258c from *Mycobacterium tuberculosis* H37Rv |
| 3d2l | 19.1 | 3.5 | 17 | Crystal structure of SAM-dependent methyltransferase (ZP_00538691.1) from *EXIGUOBACTERIUM SP. 255-15* at 1.90 A resolution |
| 5kok | 19.0 | 3.7 | 15 | Pavine N-methyltransferase in complex with Tetrahydropapaverine and S-adenosylhomocysteine pH 7.25 |
| 6uk5 | 18.9 | 3.8 | 17 | Structure of SAM bound CalS10, an amino pentose methyltransferase from *Micromonospora echinaspora* involved in calicheamicin biosynthesis |

The Dali server [27] identified numerous structures with 3D similarity to XisE (search performed on the 15th of April 2025). The best 10 hits are shown and listed according to their Z-score. Redundant protein hits have been excluded from the list.

**Table S7.** Oligonucleotides used in this study.

| Construct | Template | Primer sequence | Primer name | Reference |
|---|---|---|---|---|
| pSEVA expression plasmid linearization | pSEVA261b/ pSEVA631b | GTCGTGACTGGGAAAACCCT | AR900 | [28] |
| | | CTAGTATTTCCCCTCTTTCTCTAGT | AR970 | [28] |
| pSEVA261b-*xisA-E* | *X. hominickii* gDNA | TGTTTAATACTAGAGAAAGAGGGGAAATAC TAGATGAATAAAGATAATTTTTTACATTCT | AR1505 | This work |
| | | CTGATATATCAAGACGCGAAAAATTGGTC | AR1506 | This work |
| | *X. hominickii* gDNA | GACCAATTTTTCGCGTCTTGATATATCAG | AR1507 | This work |
| | | TCGCCAGGGTTTTCCCAGTCACGACCTTAA CTAGTAGGTTCTATGCGAGCT | AR1508 | This work |
| pSEVA261b-*xisA-C* | *X. hominickii* gDNA | TGTTTAATACTAGAGAAAGAGGGGAAATAC TAGATGAATAAAGATAATTTTTTACATTCT | AR1925 | This work |
| | | CTGATATATCAAGACGCGAAAAATTGGTC | AR1926 | This work |
| | *X. hominickii* gDNA | GACCAATTTTTCGCGTCTTGATATATCAG | AR1927 | This work |
| | | TCGCCAGGGTTTTCCCAGTCACGACTTATT TGCCTAATGTTTAATTGCC | AR1928 | This work |
| pSEVA631b-*xisDE* | *X. hominickii* gDNA | TGTTTAATACTAGAGAAAGAGGGGAAATAC TAGATGACAGTTAGAAAAATAATAGAGATT | AR1929 | This work |
| | | AGGGTTTTCCCAGTCACGACCTATGCGAG CTCTGTTTC | AR1930 | This work |
| pSEVA631b 631b-*xisE* | *X. hominickii* gDNA | TGTTTAATACTAGAGAAAGAGGGGAAATAC TAGATGAATACAGAAATATTAAAGGACTTT | AR1931 | This work |
| | | AGGGTTTTCCCAGTCACGACCTATGCGAG CTCTGTTTC | AR1930 | This work |
| pSEVA631b 631b-*xisD* | *X. hominickii* gDNA | TGTTTAATACTAGAGAAAGAGGGGAAATAC TAGATGACAGTTAGAAAAATAATAGAGATT | AR1929 | This work |
| | | GGTTTTCCCAGTCACGACCCTTTAATATTT CTGTATTCATTGTTTG | AR1932 | This work |
| pSEVA631b 261b-*xisA-C (TD::TE(xtpS))* | pSEVA261b-*xisA-C* | ACCGGATAAGGCGCAGCGGTCGGACTGAA C | AR2036 | This work |
| | | GCCCGCCTCAGCCTTAAGTATCTCT | AR2033 | This work |
| | *xtpS-TE (X. nematophila gDNA)* | AGAGATACTTAAGGCTGAGGCGGGCTATG CGGCTCCGCAGG | AR2039 | This work |
| | | TTTCCAAAAACTCCACCAAATTACTCATAAA CAGCGCCTCCACTTCGCAAT | AR2040 | This work |
| | pSEVA261b-*xisA-C* | TTTATGAGTAATTTGGTGGAGTTTTTGGAA A | AR2034 | This work |
| | | GTTCAGTCCGACCGCTGCGCCTTATCCGG T | AR2035 | This work |
| Verification for pCEP_Kan_*xisA* insertion | pCEP_Kan_*xisA* | GCTATGCCATAGCATTTTTATCCATAAG | V_pCEP_fw | This work |
| | | GCGAGTAAACCAGTAAGCATTCC | SW464 | This work |
| pEB17_Km_Δ*xisE* | *X. hominickii* gDNA | CCTCTAGAGTCGACCTGCAG_CGCAGCTT AGCCAATTAACTC | MW473 | This work |
| | | CTTCTTAACTAGTAGGTTCTATGC_CATTGT TTGAATTCCTAACACTTC | MW474 | This work |
| | *X. hominickii* gDNA | GAAGTGTTAGGAATTCAAACAATG_GCATA GAACCTACTAGTTAAGAAG | MW475 | This work |
| | | TCCCGGGAGAGCTCAGATCT_CGTATCGA CTCAATGCTTGC | MW476 | This work |
| Verification of *X. hominickii* Δ*xisE* | *X. hominickii* gDNA | CGCTTTGAACTGAGTTCGG | MW483 | This work |
| | | GGCAAATAAGTCCTTTATGACG | MW484 | This work |

| | | | | |
|---|---|---|---|---|
| pEB17_Km_ΔxisD | *X. hominickii* gDNA | CCTCTAGAGTCGACCTGCAG_TTGCGAAC GAAGCAGAAGC | MW469 | This work |
| | | CGCTTTAGTGGAGATAGGTAATC_TCTAAC TGTCATTTATTTGCCTAATG | MW470 | This work |
| | *X. hominickii* gDNA | CATTAGGCAAATAAATGACAGTTAGA_GAT TACCTATCTCCACTAAAGC | MW471 | This work |
| | | TCCCGGGAGAGCTCAGATCT_CGGGATGC GTTCATATAGC | MW472 | This work |
| Verification of *X. hominickii ΔxisD* | *X. hominickii* gDNA | CGATGAGCTGGTGATGATG | MW481 | This work |
| | | GCTGCAACCTACCGTCG | MW482 | This work |
| pCJ1 | pMLB82 | TTCAGAAAATCACGATACATGGTTTCCTTG GAGCATTAGTTG | MLB350 | This work |
| | | CCTGAAGCAACTTCACCACACAGCGCAAG TCAATCCAGTC | MLB351 | This work |
| | *V. tubiashii* gDNA | ACTAATGCTCCAAGGAAACCATGTATCGTG ATTTTCTGAACTG | MLB352 | This work |
| | | GGCACATTATGCAGGTCGACATTCGGTGT GTACTGCCAAC | MLB353 | This work |
| | pMP400 | GTTGGCAGTACACACCGAATGTCGACCTG CATAATGTGCC | MLB354 | This work |
| | | CTGTTGATCACTGATGACATCGGAATCCTC CGAATTCCTC | MLB355 | This work |
| | *V. tubiashii* gDNA | GAGGAATTCGGAGGATTCCGATGTCATCA GTGATCAACAG | MLB356 | This work |
| | | GACTGGATTGACTTGCGCTGTGTGGTGAA GTTGCTTCAGG | MLB357 | This work |
| | *V. tubiashii* gDNA | CGTGGTAATGGCGTTGAACG | MLB358 | This work |
| | | GGGACCACCATGTACAGTCG | MLB359 | This work |
| pCJ2 | pCJ1 | TGTGTTGGTCTAGCTGCCAATATGATTGGC CACAGTGAAGCTGTG | CJ23 | This work |
| | | CTGTTGATCACTGATGACATCGGAATCCTC CGAATTCCTC | MLB355 | This work |
| | pCJ1 | TTGTCGGCATTGAAAGTCCC | CJ10 | This work |
| | | GAGGAATTCGGAGGATTCCGATGTCATCA GTGATCAACAG | MLB356 | This work |
| | pCJ1 | CATATTGGCAGCTAGACCAACACAATTGTC GGTATCGTAAAGGGTG | CJ22 | This work |
| | | TGCTAGAGCAAGCTATGGCG | CJ11 | This work |

**Table S8.** Strains used in this study.

| Number | **Name** | **Description** | **Reference** |
|---|---|---|---|
| | *X. hominickii* WT | *X. hominickii* DSM 17903 WT | [29] |
| | *X. hominickii* Δ*hfq* | *hfq* is deleted | [30] |
| MW696 | *X. hominickii* $P_{BAD}$_*xisA* | pCEP_kan_*xisA* is integrated in front of *xisA* | This work |
| MW752 | *X. hominickii* Δ*xisE* $P_{BAD}$_*xisA* | *xisE* is deleted, pCEP_kan_*xisA* is integrated in front of *xisA* | This work |
| MW754 | *X. hominickii* Δ*xisD* $P_{BAD}$_*xisA* | *xisD* is deleted, pCEP_kan_*xisA* is integrated in front of *xisA* | This work |
| JC97 | *X. hominickii* Δ*hfq* $P_{BAD}$_*xisA* | *hfq* is deleted, pCEP_kan_*xisA* is integrated in front of *xisA* | This work |
| MW753 | *X. hominickii* Δ*hfq* Δ*xisE* $P_{BAD}$_*xisA* | *hfq* is deleted, *xisE* is deleted, pCEP_kan_*xisA* is integrated in front of *xisA* | This work |
| | *E. coli* ST18 | *E. coli* S17-1 *λpir ΔhemA* | [31] |
| SW189 | *E. coli* ST18 + pCEP_kan_*xisA* | *E. coli* ST18, contains pCEP_SW456/7 | This work |
| MW706 | *E. coli* ST18 + pEB17_ΔxisE | *E. coli* ST18, contains pEB17_ΔxisE | This work |
| MW708 | *E. coli* ST18 + pEB17_ΔxisD | *E. coli* ST18, contains pEB17_ΔxisD | This work |
| | *E. coli* DH10B::*mtaA* | *E. coli* DH10B with Δ*entD*::*mtaA* | [32] |
| AR 22.501 | *E. coli* DH10B::*mtaA* +*pSEVA261b* | *E. coli* DH10B::*mtaA* transformed with pSEVA261b | [28] |
| AR 22.502 | *E. coli* DH10B::*mtaA* +*pSEVA631b* | *E. coli* DH10B::*mtaA* transformed with pSEVA631b | [28] |
| AR 22.503 | *E. coli* DH10B::*mtaA* +*pSEVA261b-xisA-E* | *E. coli* DH10B::*mtaA* transformed with pSEVA261b-*xisA.E* | This work |
| AR 22.504 | *E. coli* DH10B::*mtaA* +*pSEVA261b-xisA-C* + *pSEVA631b* | *E. coli* DH10B::*mtaA* transformed with pSEVA261b-*xisA-C* and pSEVA631b | This work |
| AR 22.505 | *E. coli* DH10B::*mtaA* +*pSEVA261b-xisA-C* + *pSEVA631b-xisDE* | *E. coli* DH10B::*mtaA* *transformed with* pSEVA261b-*xisA-C* and *p*SEVA631b-*xisDE* | This work |
| AR 22.506 | *E. coli* DH10B::*mtaA* +*pSEVA261b-xisA-C* + *pSEVA631b-xisD* | *E. coli* DH10B::*mtaA* transformed with pSEVA261b-*xisA-C* and pSEVA631b-*xisD* | This work |
| AR 22.507 | *E. coli* DH10B::*mtaA* +*pSEVA261b-xisA-C* + *pSEVA631b-xisE* | *E. coli* DH10B::*mtaA* transformed with pSEVA261b-*xisA-C* and pSEVA631b-*xisE* | This work |
| AR 22.508 | *E. coli* DH10B::*mtaA* +*pSEVA261b-xisA-C (TD::TE(xtpS))* + *pSEVA631b-xisDE* | *E. coli* DH10B::*mtaA* transformed with pSEVA261b-*xisA-C (TD::TE(xtpS))* and pSEVA631b-*xisDE* | This work |
| AR 22.509 | *E. coli* DH10B::*mtaA* + *pJC1* | *E. coli* DH10B::*mtaA* transformed with pJC1 | This work |
| AR 22.510 | *E. coli* DH10B::*mtaA* + *pJC2* | *E. coli* DH10B::*mtaA* transformed with pJC2 | This work |
| | *V. cholerae* Δdef1-Δdef2 | | [3] |

**Table S9.** Plasmids used in this study.

| Plasmid | Description | Reference |
|---|---|---|
| pCEP_kan | pDS132 based, R6K ori, oriT, $Km^R$, araC, $P_{BAD}$ | [33] |
| pCEP_kan_SW456/7 | pCEP_kan with first 1046 bp of *xisA* | This work |
| pEB17 | pDS132 based, R6K ori, oriT, $Km^R$, cipB derivative with additional BglII site, sacB | [34] |
| pEB17_Km_Δ*xisE* | R6K ori, oriT, sacB, $Kan^R$ with 1098 bp upstream of *xisE* (including the first 3 nt of *xisE*) and 1063 bp downstream of *xisE* (including the last 6 nt of *xisE*) | This work |
| pEB17_Km_Δ*xisD* | R6K ori, oriT, sacB, $Kan^R$ with 1010 bp upstream of *xisD* (including the first 12 nt of *xisD*) and 1093 bp downstream of *xisD* (including the last 75 nt of *xisD*) | This work |
| pSEVA261b | pSEVA261 $araC^{AM}$ PBad RiboJ B0064-RBS | [28] |
| pSEVA631b | pSEVA631 $araC^{AM}$ PBad RiboJ B0064-RBS | [28] |
| pSEVA261b-*xisA-E* | *xisA-E* inserted under the control of $P_{BAD}$ | This work |
| pSEVA261b-*xisA-C* | *xisA-C* inserted under the control of $P_{BAD}$ | This work |
| pSEVA631b-*xisDE* | *xisA-C* inserted under the control of $P_{BAD}$ | This work |
| pSEVA631b-*xisD* | *xisA-C* inserted under the control of $P_{BAD}$ | This work |
| pSEVA631b-*xisE* | *xisA-C* inserted under the control of $P_{BAD}$ | This work |
| +pSEVA261b-*xisA-C* (TR::TE(xtpS)) + pSEVA631b-*xisDE* | *xisA-C* inserted under the control of $P_{BAD}$; XisB TR domain was exchanged by the TE domain from XtpS from *X. nematophila* | This work |
| pMLB82 | | [3] |
| pMLB82 | | [3] |
| pJC1 | *oriV*(RK2); Δ*xylS*-Pm expression system; KmR; $oriT_{RP4}$; $P_{def2VCH}$-PPTASE; $P_{BAD}$-VtBGC-$Vtdef_{BGC}$+ | This work |
| pCJ2 | *oriV*(RK2); Δ*xylS*-Pm expression system; KmR; $oriT_{RP4}$; $P_{def2VCH}$-PPTASE; $P_{BAD}$-VtBGC-$Vtdef_{BGC}$- | This work |

### D. Supplementary Figures

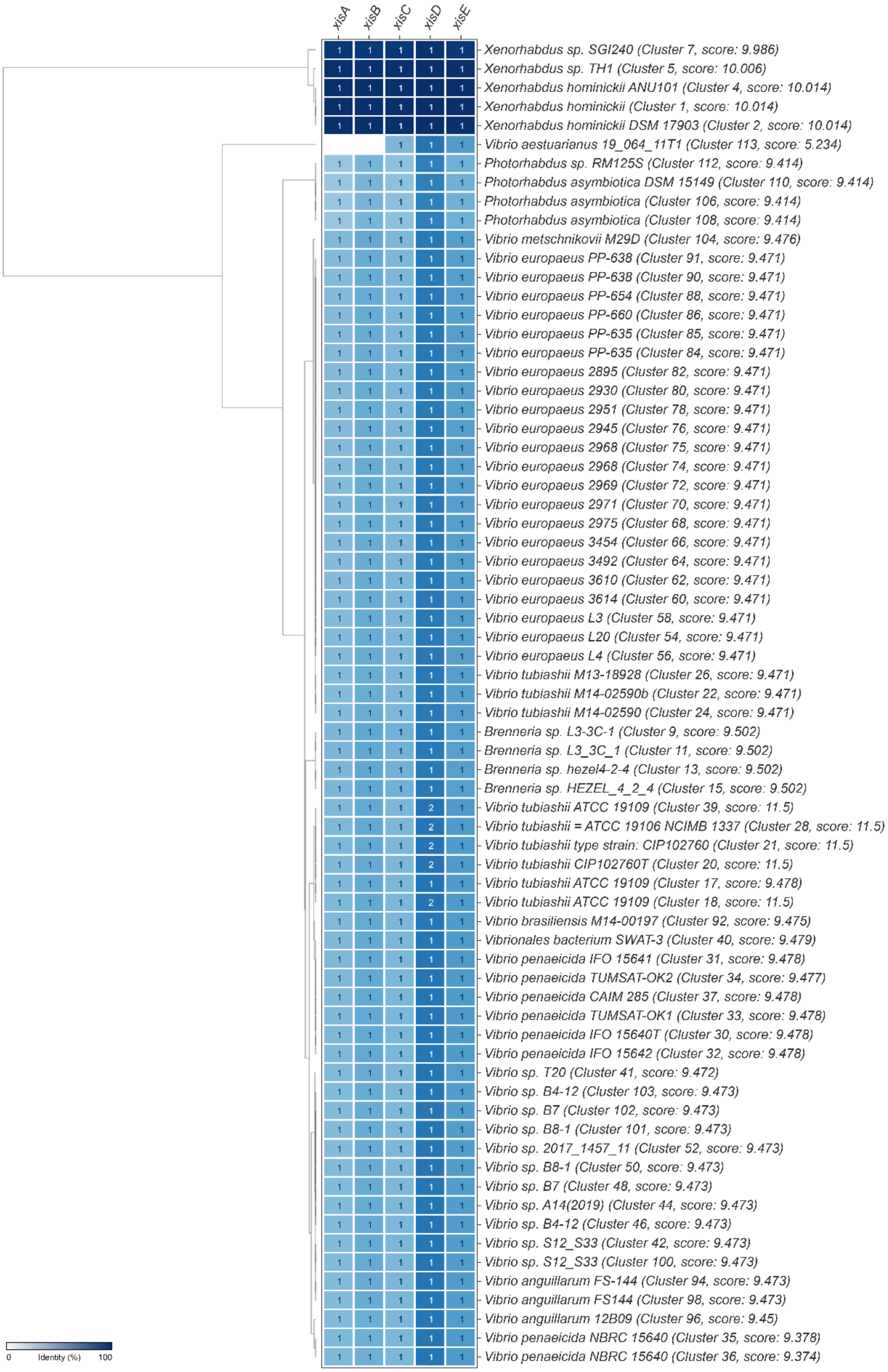


**Figure S1.** Complete CAGECAT analysis for the *xis* BGC. As query file, *xisA-E* from *X. hominickii* was selected.

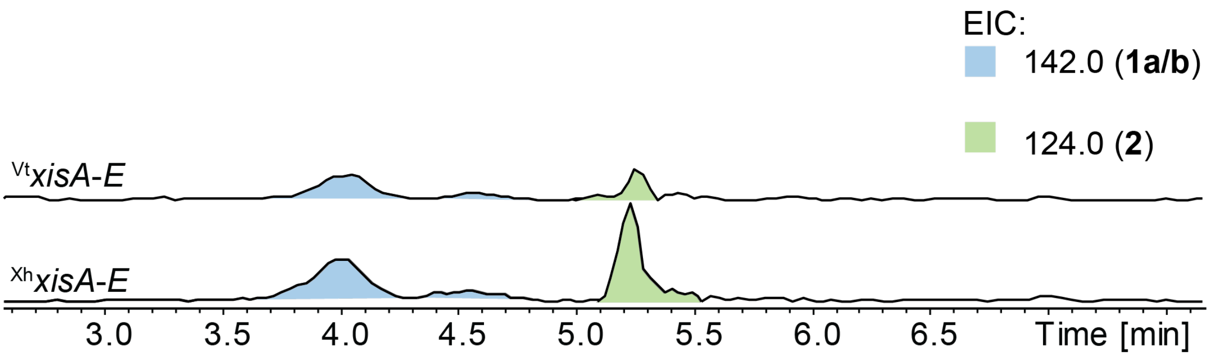


**Figure S2.** EICs of Xildivaline production from heterologous expression of the *Vibrio tubiashii* (Vt) BGC and *X. hominickii* (Xh) in *E. coli mtaA*.

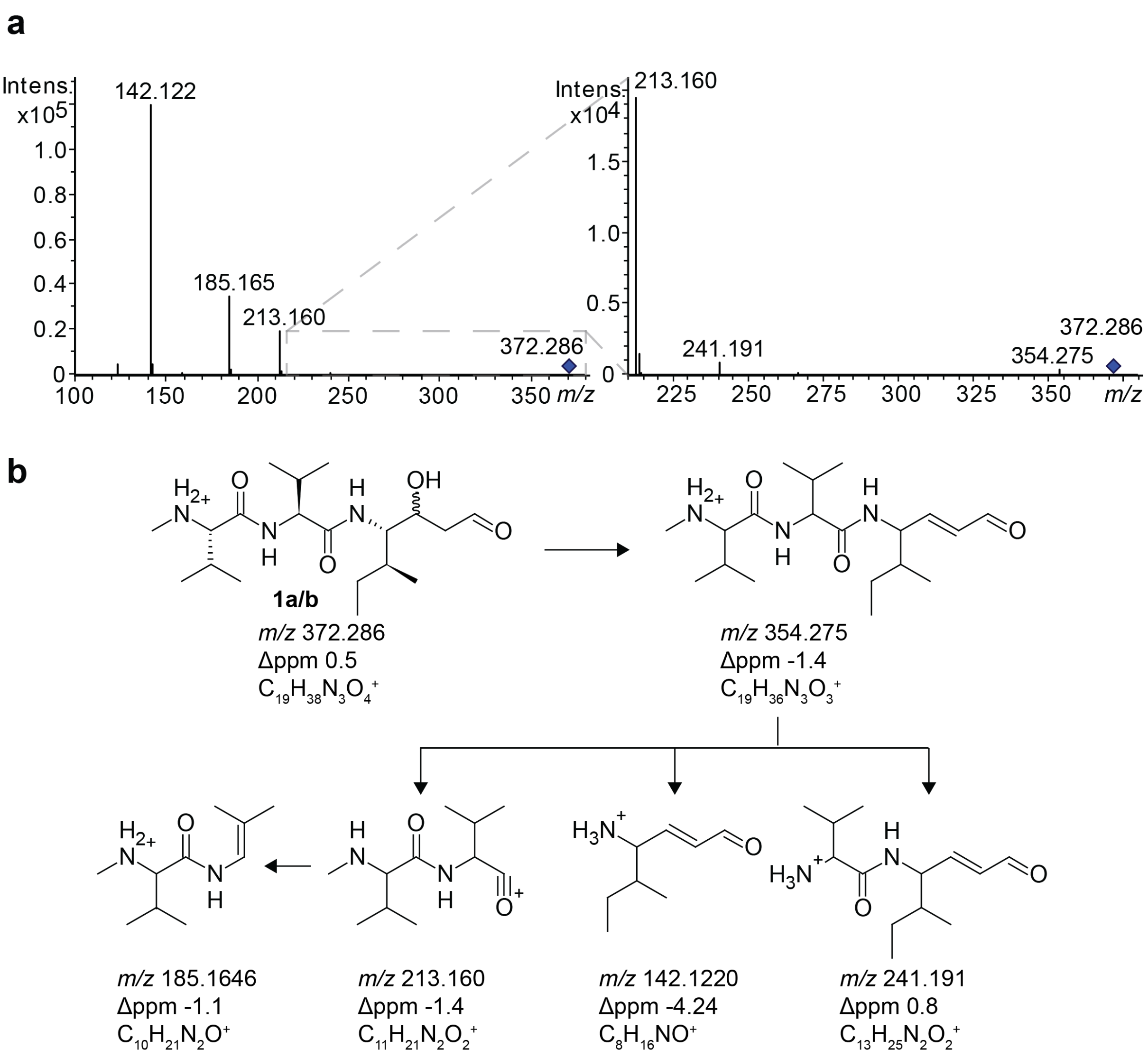


**Figure S3.** MS/MS spectrum (**a**) and observed fragments (**b**) of **1a/b**.

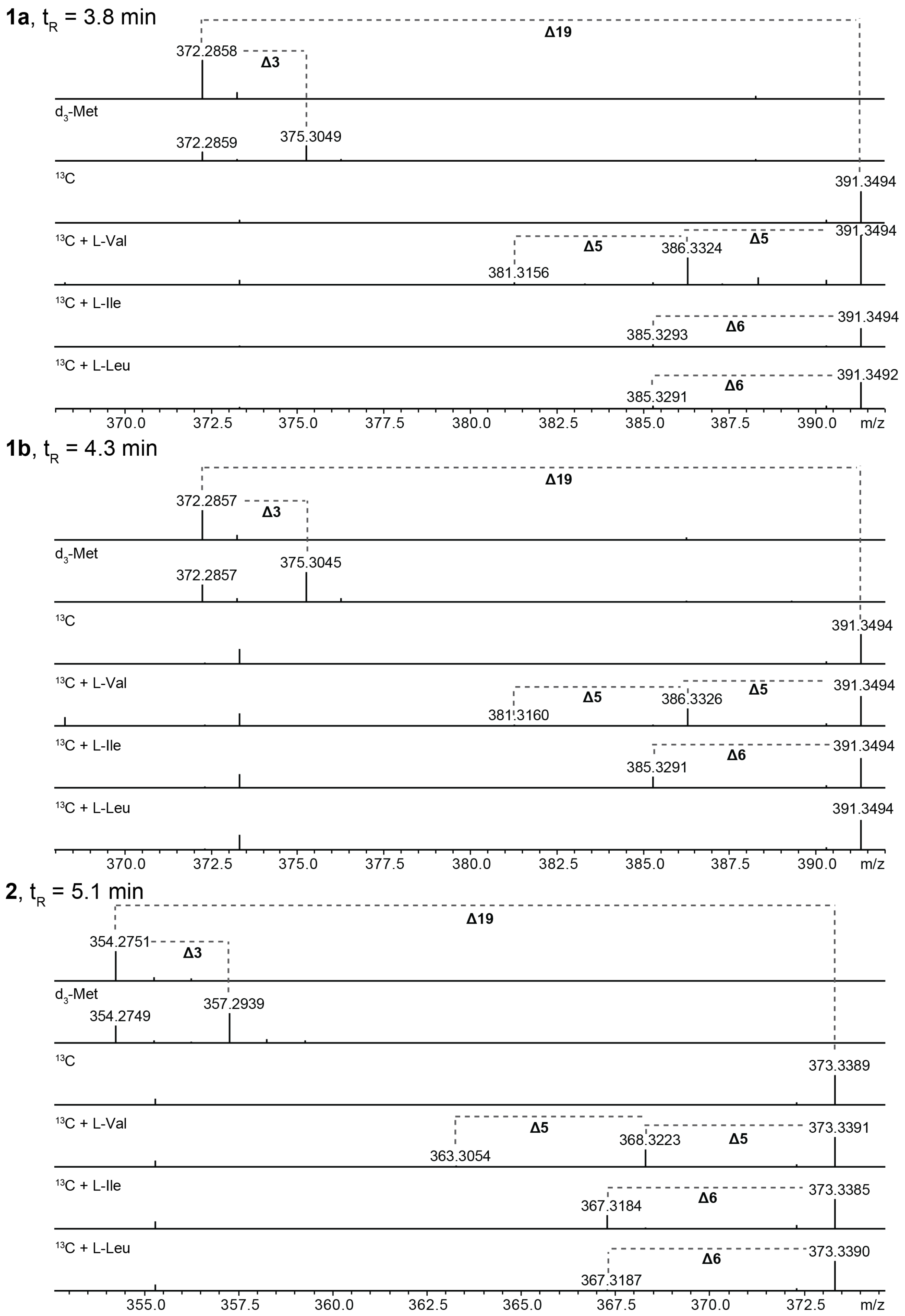
1a, tR = 3.8 min
Δ19
372.2858
Δ3
d3-Met
372.2859
375.3049
13C
391.3494
13C + L-Val
391.3494
Δ5
386.3324
Δ5
381.3156
13C + L-Ile
391.3494
Δ6
385.3293
13C + L-Leu
391.3492
Δ6
385.3291
370.0
372.5
375.0
377.5
380.0
382.5
385.0
387.5
390.0
m/z
1b, tR = 4.3 min
Δ19
372.2857
Δ3
d3-Met
375.3045
372.2857
13C
391.3494
13C + L-Val
Δ5
386.3326
Δ5
391.3494
381.3160
13C + L-Ile
391.3494
Δ6
385.3291
13C + L-Leu
391.3494
370.0
372.5
375.0
377.5
380.0
382.5
385.0
387.5
390.0
m/z
2, tR = 5.1 min
Δ19
354.2751
Δ3
d3-Met
354.2749
357.2939
13C
373.3389
13C + L-Val
Δ5
368.3223
Δ5
373.3391
363.3054
13C + L-Ile
373.3385
Δ6
367.3184
13C + L-Leu
373.3390
Δ6
367.3187
355.0
357.5
360.0
362.5
365.0
367.5
370.0
372.5
m/z

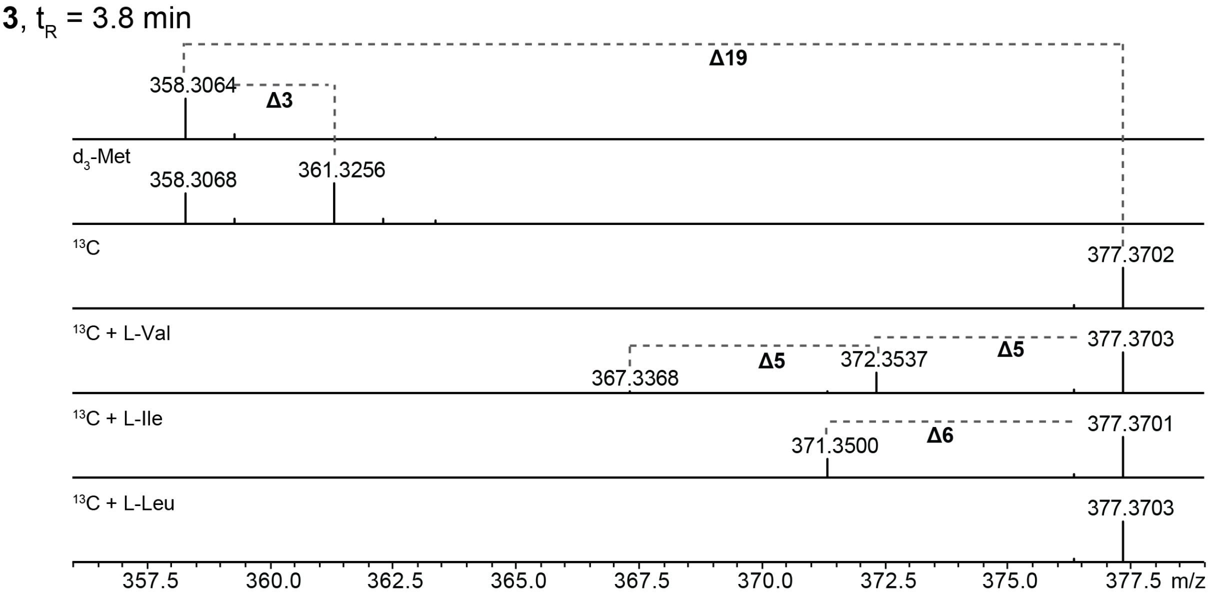


**Figure S4.** MS data of xildivaline derivatives **1-3** cultivated in different growth media using *methyl*-[$D_3$]methionine in XPPM and non-labeled amino acids in fully $^{13}$C labelled ISOGRO® medium.

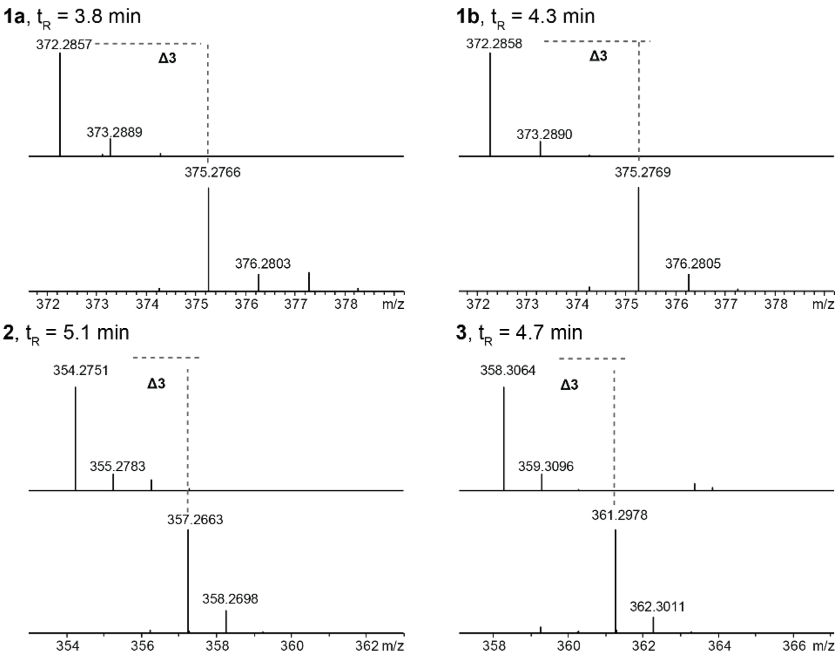


**Figure S5.** MS data of xildivaline derivatives **1-3** from cultures grown in LB ($^{14}$N, top) and ISOGRO®-$^{15}$N medium (bottom).

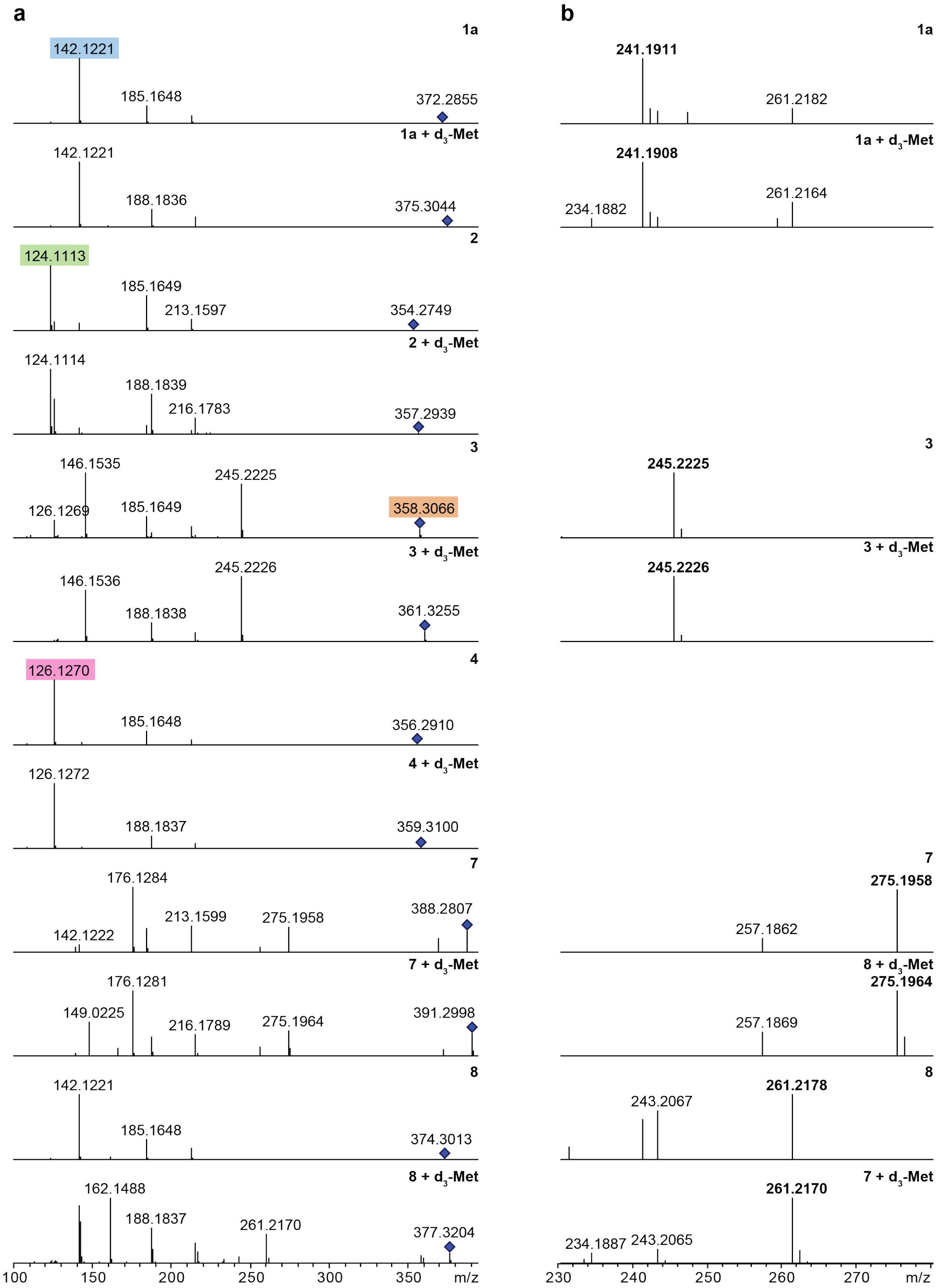
a
1a
142.1221
185.1648
372.2855
1a + $d_3$-Met
142.1221
188.1836
375.3044
2
124.1113
185.1649
213.1597
354.2749
2 + $d_3$-Met
124.1114
188.1839
216.1783
357.2939
3
146.1535
245.2225
126.1269
185.1649
358.3066
3 + $d_3$-Met
245.2226
146.1536
188.1838
361.3255
4
126.1270
185.1648
356.2910
4 + $d_3$-Met
126.1272
188.1837
359.3100
7
176.1284
213.1599
275.1958
388.2807
142.1222
7 + $d_3$-Met
176.1281
149.0225
216.1789
275.1964
391.2998
8
142.1221
185.1648
374.3013
8 + $d_3$-Met
162.1488
188.1837
261.2170
377.3204
100
150
200
250
300
350
m/z
b
1a
241.1911
261.2182
1a + $d_3$-Met
241.1908
234.1882
261.2164
3
245.2225
3 + $d_3$-Met
245.2226
7
275.1958
257.1862
8 + $d_3$-Met
275.1964
257.1869
8
261.2178
243.2067
7 + $d_3$-Met
261.2170
234.1887
243.2065
230
240
250
260
270
m/z

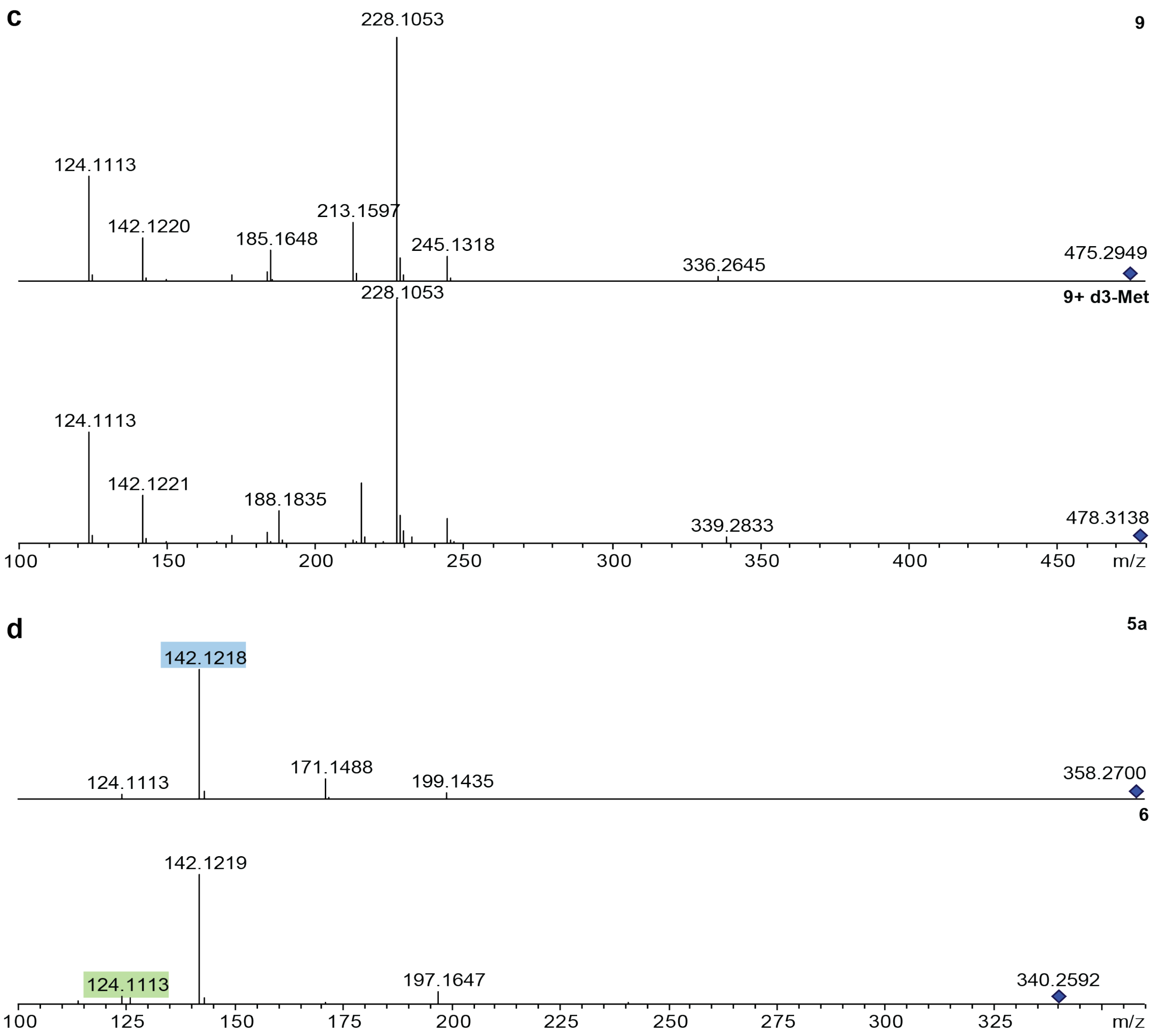


**Figure S6.** MS/MS spectra of selected xildivaline derivatives **1a** - **9**. MS/MS spectra data of xildivaline derivatives **1a** - **8** cultivated in XPPM with and without the addition of *methyl*-[$D_3$]methionine (**a**). Zoomed range of *m/z* 230 - 280 for a better visualization of the "valyl-isoleucyl-polyketide" fragment ion (highlighted in bold), no corresponding masses could be detected for **3** and **8** (**b**). MS/MS spectra data of xildivaline derivatives **9** (**c**), **4a** and **5** (**d**) cultivated in XPPM with and without the addition of *methyl*-[$D_3$]methionine.

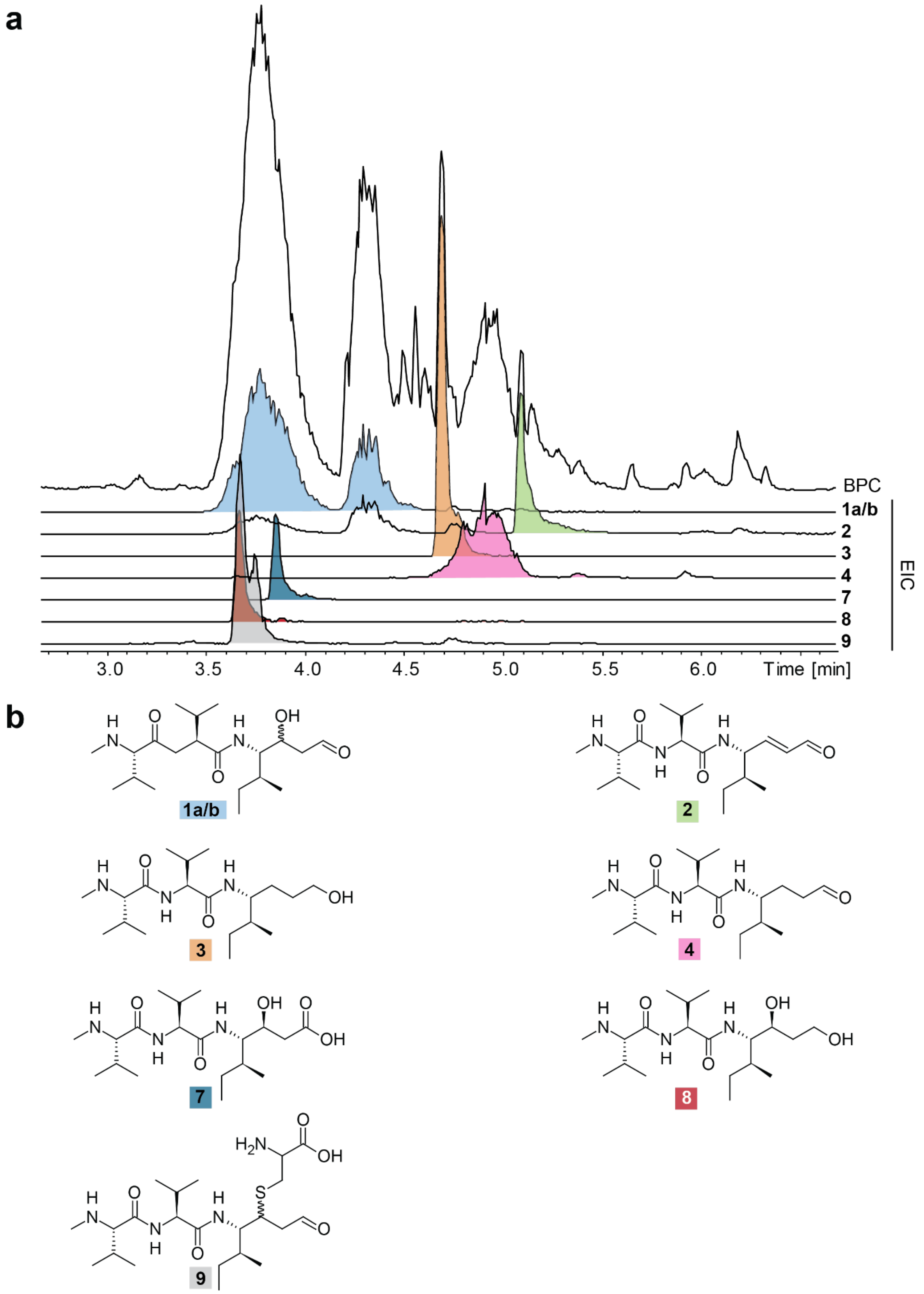


**Figure S7.** HPLC/MS chromatogram of concentrated extract from *X.hominickii* WT P-$_{BAD}$-xisA showing minor amounts of **7-9**. No non-methylated derivatives like **5** were detected.

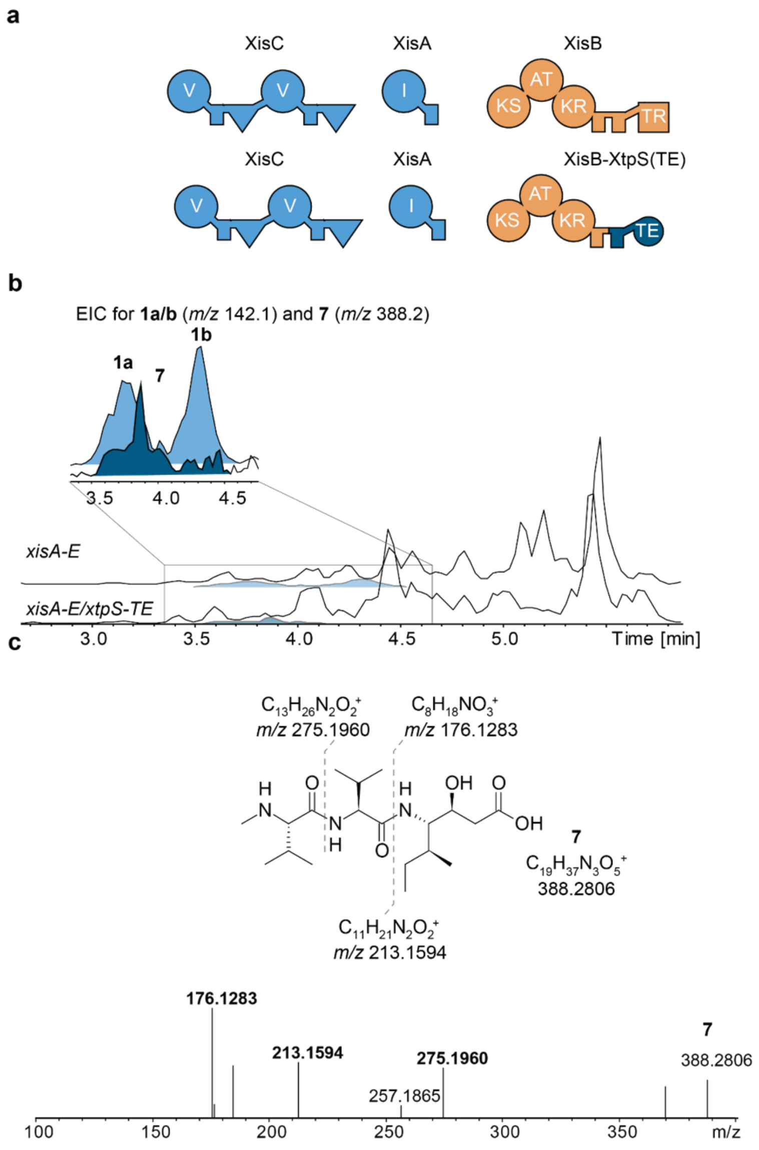


**Figure S8.** Comparison between natural and engineered XisB variant showing replacement of the terminal T-TR against a T-TE domain from the xenotetrapeptide producing NRPS XtpS (**a**), resulting in the production of **7** carrying a C-terminal carboxylic acid (**b**), also confirmed by MS/MS analysis (**c**).

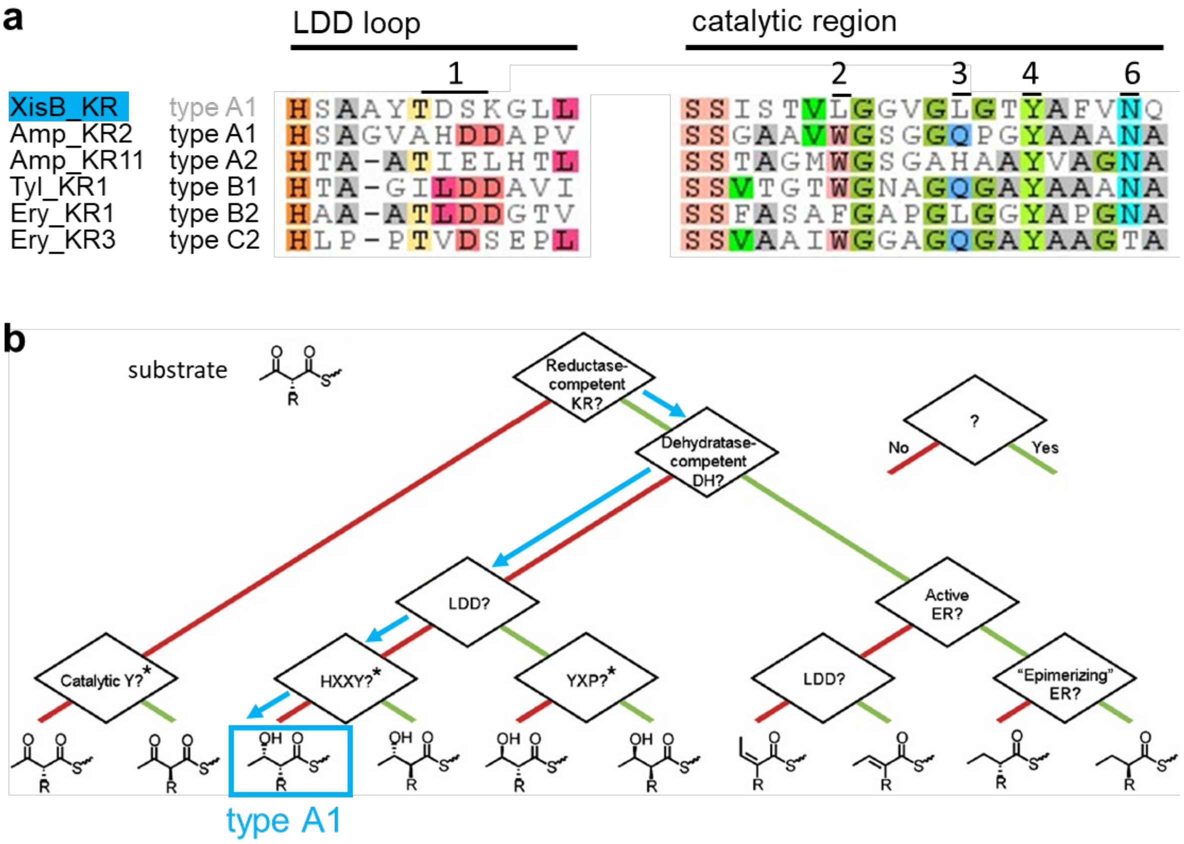


**Figure S9.** XisB_KR in silico analysis based on the protocol by Keatinge-Clay.[35] The two described ketoreductases allow the substrate to enter the active site either from a the left side via a conserved tryptophan (for XisB_KR a Leu) (type A) or the right side via a LDD motif (type B) determining the stereo outcome as S (type A) or R (type B). (**a**) LDD loop and catalytic region are aligned with representatives from five different KR types. XisB_KR shows no LDD motif (1) present in B type ketoreductases and shows leucine (2) instead of conserved tryptophan found in A type ketoreductases as also found in other type A KRs. [36] XisB_KR shows no H in position 3 and the catalytic tyrosine in position 4. (**b**) Subsituent flowchart after Keatinge-Clay determines XisB_KR as type A1 KR. [35] XisB exhibits no DH domain, XisB_KR shows no LDD motif (1) and no HXXY motif (3-4) thus suggesting S configuration of the hydroxy group by this A type KR.

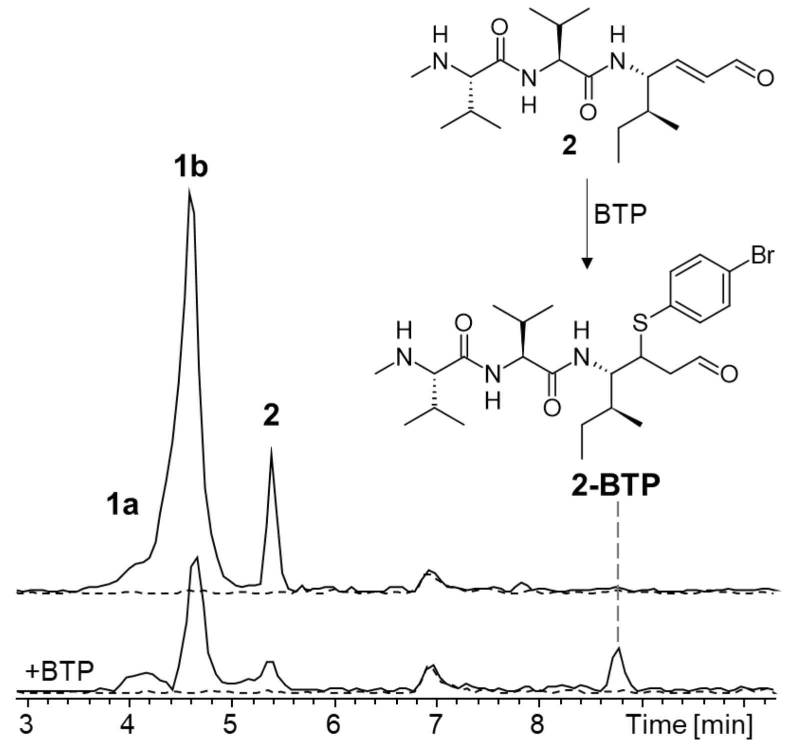


**Figure S10.** 4-Bromthiophenol (BTP) addition to **2** confirms its Michael acceptor leading to the BTP adduct **2-BTP** with a *m/z* 542.2. The *m/z* of the MeVal-Val fragment 213.1 are shown as control. Induced BPCs are indicated by solid lines and non-induced cultures by dashed lines.

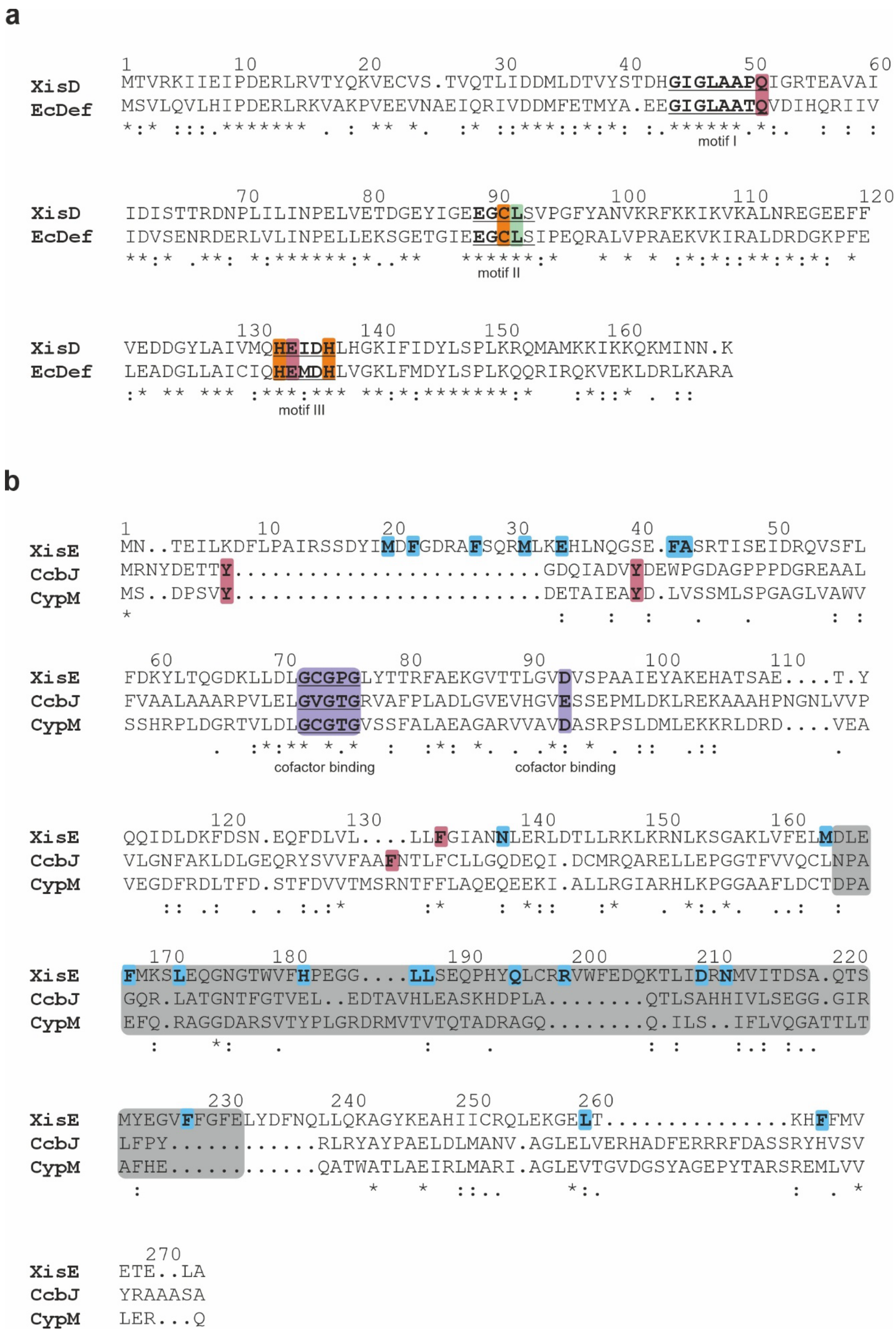


**Figure S11. Sequence comparison of (a) XisD and (b) XisE with their structural relatives.** (**a**) Sequence alignment of XisD with *Escherichia coli* deformylase (EcDef). Motifs characteristic for deformylases (GϕGϕAAXQ, EGCϕS, and HEϕDH with ϕ being a hydrophobic and X any amino acid) are underlined and labelled. Residue numbers are indicated for XisD. Amino acids coordinating the metal ion in the active site of XisD and EcDef are highlighted in orange. Residues involved in formiate binding are red (interactions via side chains) and green (interaction via main chain atoms). (**b**) Sequence alignment of XisE with CcbJ from *Streptomyces caelestis* and CypM from *Streptomyces sp*. Residue numbers are provided for XisE. The GxGxG sequence motif as well as the acidic residue at the end of β-sheet 2, typical of type I methyltransferases and relevant for cofactor binding within the Rossmann domain, are shaded purple. The substrate binding domain of XisE, CcbJ and CypM is colored gray. Amino acids relevant for the catalytic activity of CcbJ (highlighted in red) are absent from XisE. Residues forming the substrate binding pocket of XisE are highlighted against a blue background. Phe133 (red) might be catalytically relevant.

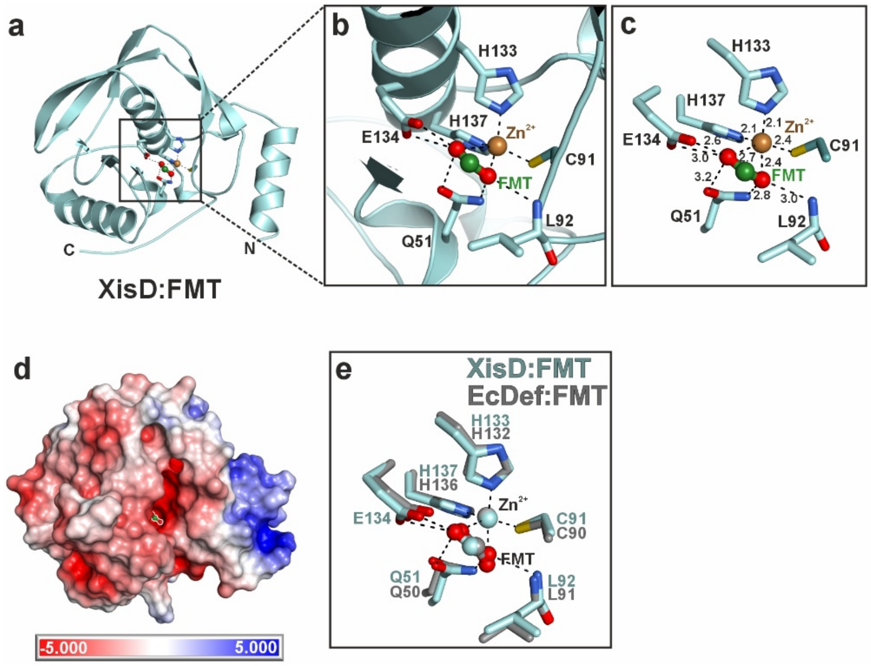


**Figure S12. X-ray structure of the deformylase XisD in complex with formiate (FMT).** (**a**) Ribbon illustration of XisD with bound FMT. N- and C-termini are labelled. (**b**) Zoom-in at the XisD active site. Key residues coordinating the $Zn^{2+}$ ion (brown sphere) and the FMT (ball-and-stick model in red and green) are shown as sticks and labelled. Hydrogen bonds are indicated by black dotted lines. (**c**) Close-up view of the XisD active site with all distances for interactions given in Å. (**d**) Connolly surface representation of XisD. Colors indicate negative (red) and positive (blue) electrostatic surface potential, ranging from -5kT/e to + 5kT/e. FMT is shown as ball-and-stick model at the bottom of the substrate binding pocket. (**e**) Superposition of active site residues of XisD:FMT (lightblue) and *E. coli* deformylase (EcDef, PDB entry 1XEM [23]; gray), both in complex with $Zn^{2+}$. All residues for FMT and metal coordination are conserved.

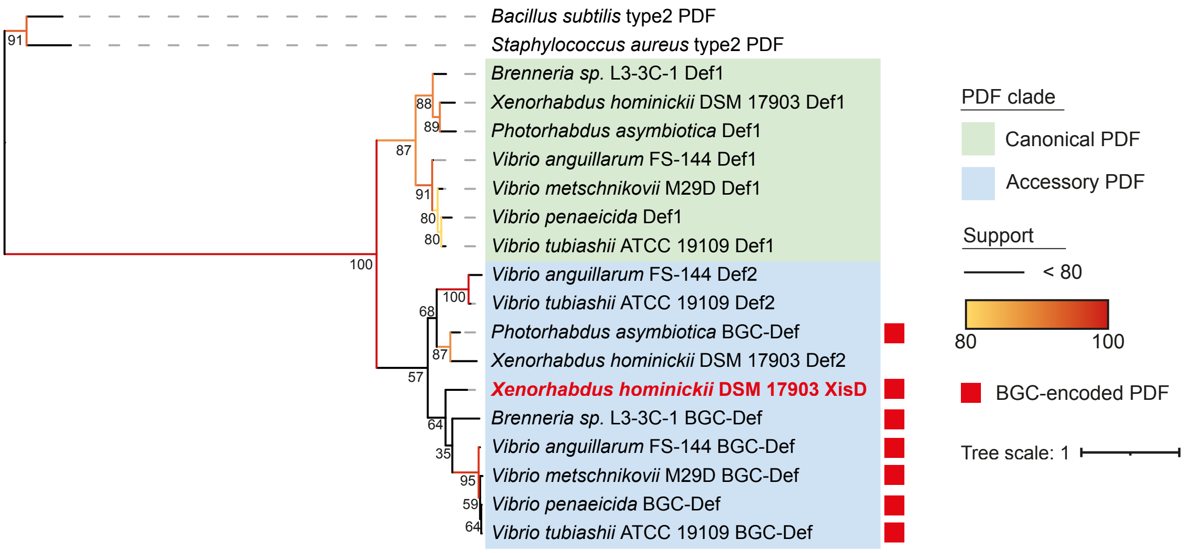


**Figure S13.** Phylogenetic tree of PDFs encoded by seven Gammaproteobacteria harboring a xildivaline-like BGC (see also Fig. S1). The BGC-encoded PDF XisD is highlighted in red. UFBoot support values >80 are shown using a yellow-to-red gradient. The tree was rooted using Def sequences from Gram-positive bacteria as outgroup.

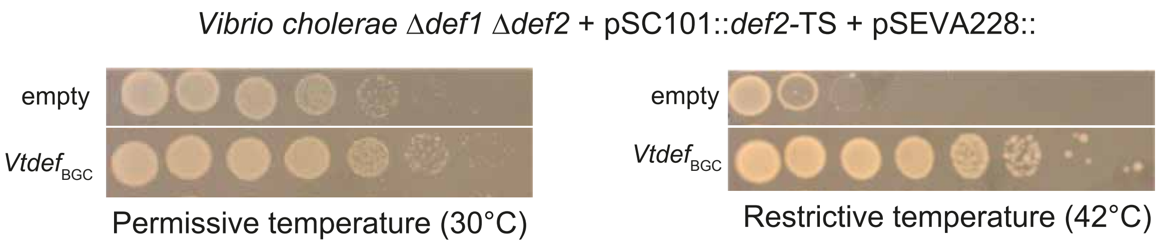


**Figure S14.** ***In vivo*** **activity of the** ***V. tubiashii*** **BGC-associated PDF.** Drop test of *V. cholerae* Δ*def1*$_{VCH}$ Δ*def2*$_{VCH}$ harboring a pSC101::*def2*$_{VCH}$-TS plasmid and a pSEVA228 plasmid at permissive temperature (left panel) or restrictive temperature (right panel). Growth at restrictive temperature indicate that the PDF coding in the pSEVA228 plasmid is enzymatically active.

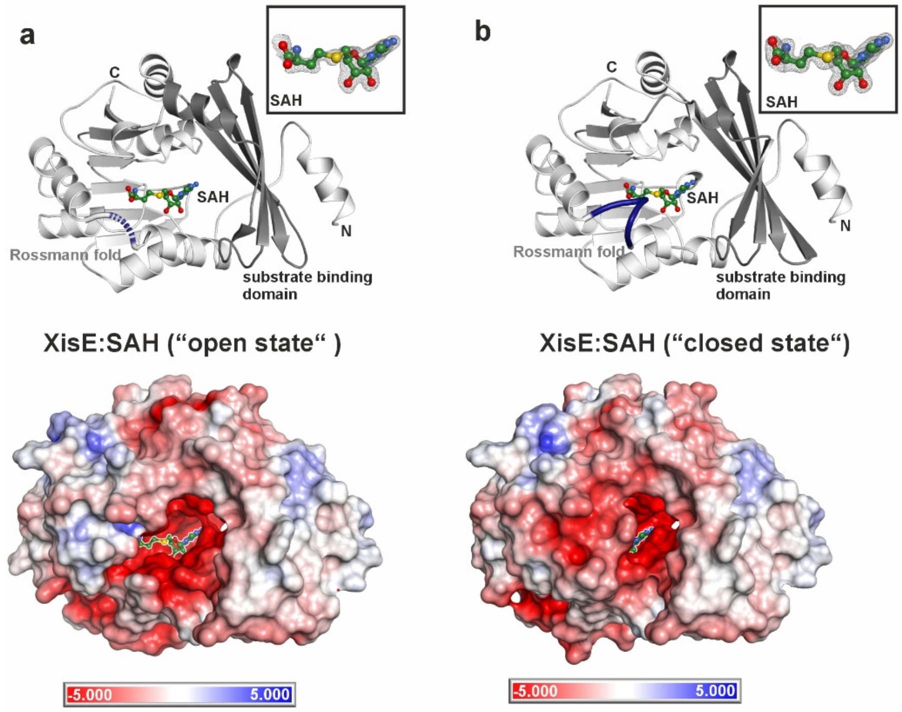


**Figure S15. X-ray structure of XisE in complex with SAH.** (**a**) Upper panel: Cartoon illustration of XisE with its substrate binding domain, consisting of a four-stranded antiparallel β-sheet, shown in dark gray. Residues 38-44 are disordered as indicated by a thickened dotted purple loop. This state is denoted as the “open state” of XisE. Although SAH (shown as a ball-and-stick model) is bound, its $2F_O$-$F_C$ omit electron density map (gray mesh contoured to 1σ) indicates reduced occupancy (box in the upper right corner). Lower panel: Negative (red) and positive (blue) electrostatic surface potential (-5kT/e to + 5kT/e) of XisE. (**b**) Ribbon structure and electrostatic surface potential of XisE in the “closed state”. Residues 38-44 (thickened purple segment) are ordered and form a substrate binding loop that shapes the wide and open active site pocket. The occupancy of SAH is higher than in the open state as confirmed by its well-defined $2F_O$-$F_C$ omit electron density map (gray mesh contoured to 1σ; boxed in the right upper corner).

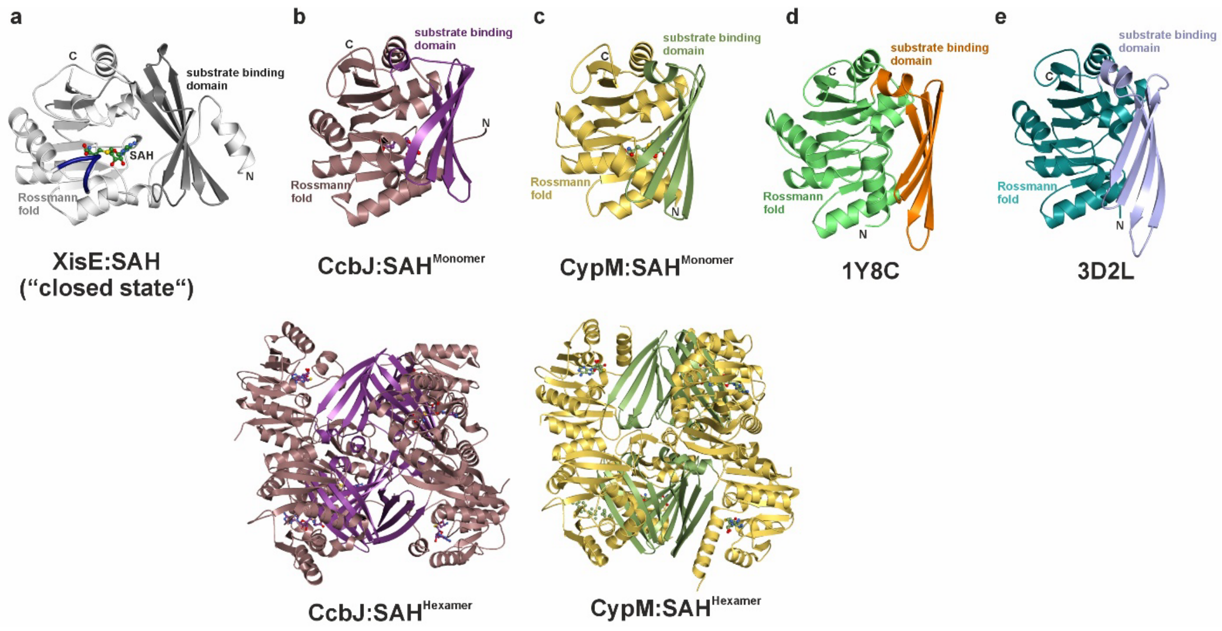


**Figure S16. Comparison of XisE with structurally related methyltransferases.** (**a**) Ribbon illustration of XisE with bound SAH in the "closed state". The substrate binding domain consisting of a four-stranded antiparallel β-sheet is colored in dark gray. (**b**) Cartoon representation of a CcbJ monomer in complex with SAH (upper panel; PDB entry 4HH4 [25]). The substrate binding domain (purple) is rotated inwards in comparison to XisE (panel a). CcbJ oligomerizes to a hexamer via the substrate binding domain (lower panel). (**c**) Ribbon structure of a CypM monomer with bound SAH (upper panel; PDB entry 7WZG, Duan, S.Y., unpublished). Like in CcbJ, the substrate binding domain is rotated towards the Rossmann fold and CcbJ assembles into hexamers (lower panel). (**d**) A methyltransferase from *Clostridium acetobutylicum* ATCC 824 (PDB entry 1Y8C, unpublished) also has its substrate binding domain rotated inwards but does not form oligomers. (**e**) Similarly, the substrate binding domain of monomeric methyltransferase (ZP_00538691.1) from *Exiguobacterium sp* (PDB entry 3D2L, unpublished) is flipped inwards.

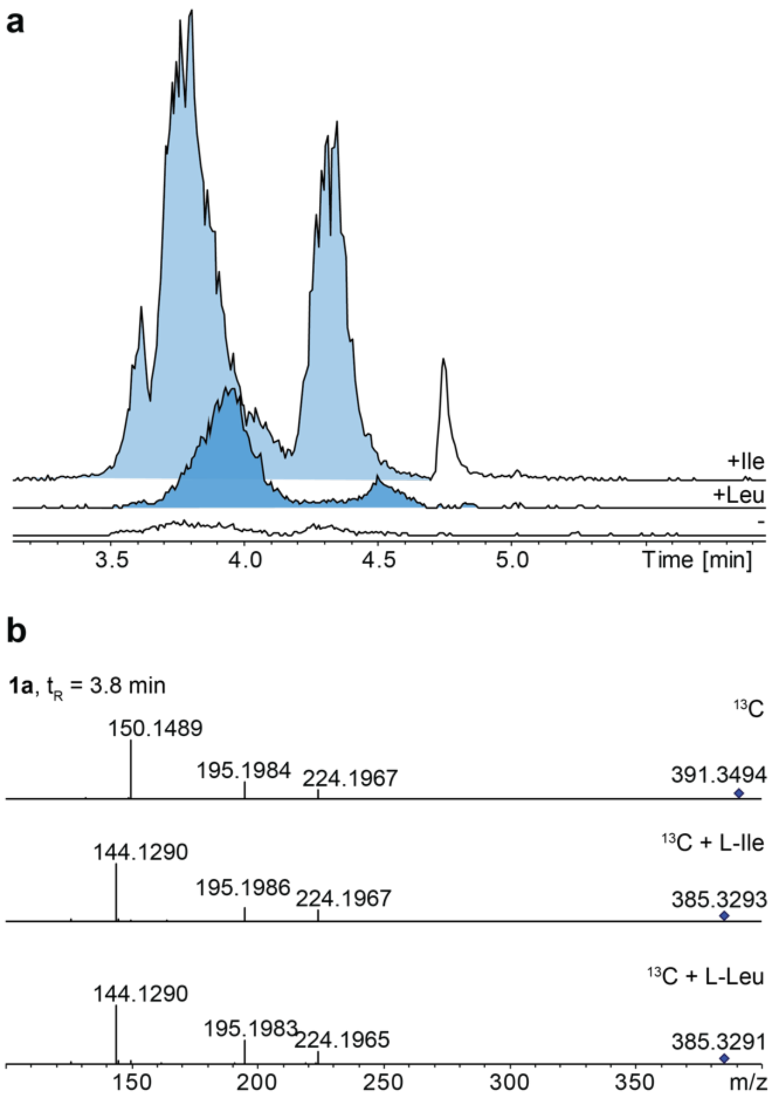


**Figure S17.** Extracted Ion Chromatograms (EICs) for *m/z* 385.329 from *X. hominickii* Δhfq $P_{BAD}$-*xisA* cultivated in ISOGRO®-$^{13}$C medium with the addition of $^{12}$C-Ile or $^{12}$C-Leu (**a**) and observed MS/MS spectra, against *m/z* 391.3494 (fully $^{13}$C labelled **1a**/**b**) (**b**).
